# Consistent crown phenology, not functional traits, predicts whether tropical trees can be mapped from imaging spectroscopy

**DOI:** 10.64898/2026.05.06.722428

**Authors:** James G C Ball, Sadiq Jaffer, Anthony Laybros, Colin Prieur, Anil Madhavapeddy, Nicolas Barbier, Grégoire Vincent, David A Coomes

## Abstract

- Airborne imaging spectroscopy can identify tree species in hyperdiverse tropical forests, but accuracy varies greatly among species. We asked which ecological and evolutionary attributes make a tropical tree species spectrally separable.
- In moist tropical forest in French Guiana, we fitted the learning curve of accuracy against the number of labelled crowns for 169 species, then tested six hypothesised determinants on 45 species rarefied to ten crowns, at species, species-pair and crown scales.
- Dispersed sampling and larger crowns increased accuracy, and liana infestation more than halved the odds of correct identification. Phenological regularity, how closely conspecific crowns share one leaf cycle, in step or not, was the strongest species-level correlate and survived every multivariate analysis. It did not act by making conspecifics uniform on the acquisition date: regular species were spectrally more distinct from other species, and within species the crowns that best followed the species’ leaf cycle were classified correctly. Congeners reduced accuracy; leaf and bark traits did not predict separability.
- Separability depends on the consistency of a species’ signal, how faithfully its crowns express a shared leaf cycle, how widely it is sampled and how free its crowns are from contamination, rather than on its measured functional traits.

## 1 Introduction

Species-level maps of tropical forests underpin a wide range of ecological and conservation applications, from tracking species abundance and health to improving carbon-stock estimates and informing restoration planning [Chave et al., 2003, Fassnacht et al., 2016, Davies et al., 2021, Weinstein et al., 2024]. Airborne hyperspectral imagery, which captures canopy reflectance across hundreds of narrow spectral bands, provides the most promising current pathway to mapping individual species in complex forests [Clark et al., 2005, Féret and Asner, 2013, Laybros et al., 2019]. Ball et al. [2026] demonstrate that combining convolutional neural network (CNN)-based multi-temporal crown segmentation with hyperspectral classification can achieve a weighted F1 of 0.75 across 169 species at the Paracou Field Station, French Guiana – a substantial advance over previous studies typically limited to fewer than 20 species. However, this aggregate accuracy masks enormous variation among species: repeated cross-validation showed that 38 species maintained mean F1 ≥ 0.7, while many remained effectively unclassifiable.

What determines whether a given species can be reliably identified from its canopy reflectance spectrum, its mappability? The question is fundamental to the limits of remote sensing for biodiversity monitoring and to the design of data collection, yet it has received little systematic attention. The number of labelled training crowns is expected to limit accuracy for rare species, which dominate tropical forests [ter Steege et al., 2013, Baldeck and Asner, 2014]; it is a property of the sample rather than of the species, and the learning curve of accuracy against it is the baseline on which every other factor must be compared. The most direct candidate is the species’ spectral distinctiveness, whether it occupies a compact, well-separated region of spectral space [Clark et al., 2005, Féret and Asner, 2013]: any other attribute can affect separability only through it. Two further factors concern how a species is sampled and how clean its crowns are. Where its crowns lie matters: tightly clustered training individuals can inflate apparent accuracy through spatial autocorrelation [Kattenborn et al., 2022], whereas individuals dispersed across varied conditions give the classifier a more representative sample and a sterner test of whether it has learned the species rather than the place [Marconi et al., 2022]. And liana infestation introduces foreign foliage into the crown spectrum, degrading the species-specific signal [Castro-Esau et al., 2004, van der Heijden et al., 2022].

The remaining factors concern the biology of the species. Tree stature and crown size should matter because a large crown contributes more pixels and a smaller share of edge pixels mixed with neighbours, understorey and shadow, and discrimination of tropical species is known to depend on the scale at which spectra are measured [Clark et al., 2005]. Leaf and bark traits influence reflectance [Asner and Martin, 2009, Ustin et al., 2009], though how far individual traits predict classification success remains unclear. Leaf phenology, the timing of leaf production, senescence and shedding, strongly modulates crown reflectance [Blackburn and Milton, 1995, Clark and Roberts, 2012], and the obvious expectation for a single-date image is that species whose individuals are in step on the acquisition date present a homogeneous signal, whereas asynchronous or weakly seasonal species vary more among themselves [Hesketh and Sánchez-Azofeifa, 2012]. Conspecifics may also share the same leaf cycle without being in step; that property is regularity, how much of a crown’s leaf cycle is species rather than individual, as distinct from synchronicity, how much of it is also season. Finally, evolutionary relatedness shapes the spectral landscape because closely related species often retain similar leaf chemistry and structure [Cavender-Bares et al., 2016, McManus et al., 2016, Meireles et al., 2020], and whether that signal acts through deep clade structure or through the nearest relatives present matters for forests of different composition.

Previous studies have explored individual aspects of spectral separability, but none that we know of has evaluated these factors together in a tropical forest, or across scales from crowns through species pairs to the community. Here we build on work at Paracou, French Guiana, where 3,256 tree crowns of 239 species were hand-delineated and verified in the field in permanent inventory plots, a single airborne hyperspectral acquisition covered the plots, and a spectrum was extracted for every crown [Ball et al., 2026]. Apart from the learning curve and the error structure of the full classification, every analysis here re-runs the classification in experiments designed for this question. The learning curve of accuracy against the number of field-labelled crowns per species, fitted to the 169 species of the reference dataset, set the design: so that differences in sample size would not drive the result, we kept the 45 species with at least ten crowns and complete predictor data and rarefied each to ten crowns in every classification run. This standardised dataset is the basis of the systematic evaluation that follows.

Our first test was whether spectrally distinct species were more easily identified. We expected this, since it is how a linear discriminant classifier assigns species, so the test serves as a check on the classifier and establishes the proximate basis through which every other factor must act. We then tested the hypotheses that more easily distinguished species (1) were more widely dispersed across the landscape, (2) were free of liana infestation, (3) were tall with broad crowns, (4) had distinctive leaf and bark traits, (5) showed strong, regular and, above all, synchronised leaf phenology, and (6) were evolutionarily and taxonomically isolated. We explored these hypotheses with two measures of separability. *One-vs-all separability*, the species’ F1 score in the multi-class classifier (hereafter simply F1), asks how well a species can be distinguished from the whole community, and was related to the candidate predictors by univariate correlations and then by elastic net and beta regression. *Pairwise separability* asks how distinct two particular species are from each other, measured as the spectral distance between them and the rate at which the classifier confuses them. Finally, a crown-level model asks which attributes of an individual crown govern its classification. By asking whether complementary methods converge on the same predictors, we aim to provide the first comprehensive, multi-scale assessment of the ecological and evolutionary determinants of spectral separability.

## 2 Materials and Methods

### 2.1 Study site and tree-crown dataset

This study builds on Ball et al. [2026], which predicted the species of individual tree crowns in hyperdiverse moist tropical forest from hyperspectral imagery. We describe that work briefly and explain how its dataset of field-labelled crowns was reduced to the 45 species, each with at least ten training crowns and complete predictor data, analysed here.

Fieldwork was conducted at the Paracou Field Station in French Guiana (5*^◦^* 16*^′^* N, 52*^◦^* 55*^′^* W), a lowland moist tropical forest described in detail in Ball et al. [2026] (Fig. 1). The site receives 3,200 mm of annual rainfall, with a dry season from mid-August to mid-November [Bonal et al., 2008, Wagner et al., 2011]. Its 27 permanent plots (0.5–25 ha) contain 76,000 trees of diameter at breast height (DBH) ≥10 cm spanning over 800 species, inventoried every 1–5 years [Gourlet-Fleury et al., 2004], 90% of which have been placed in a time-calibrated phylogeny [Baraloto et al., 2012].

**Figure 1:**
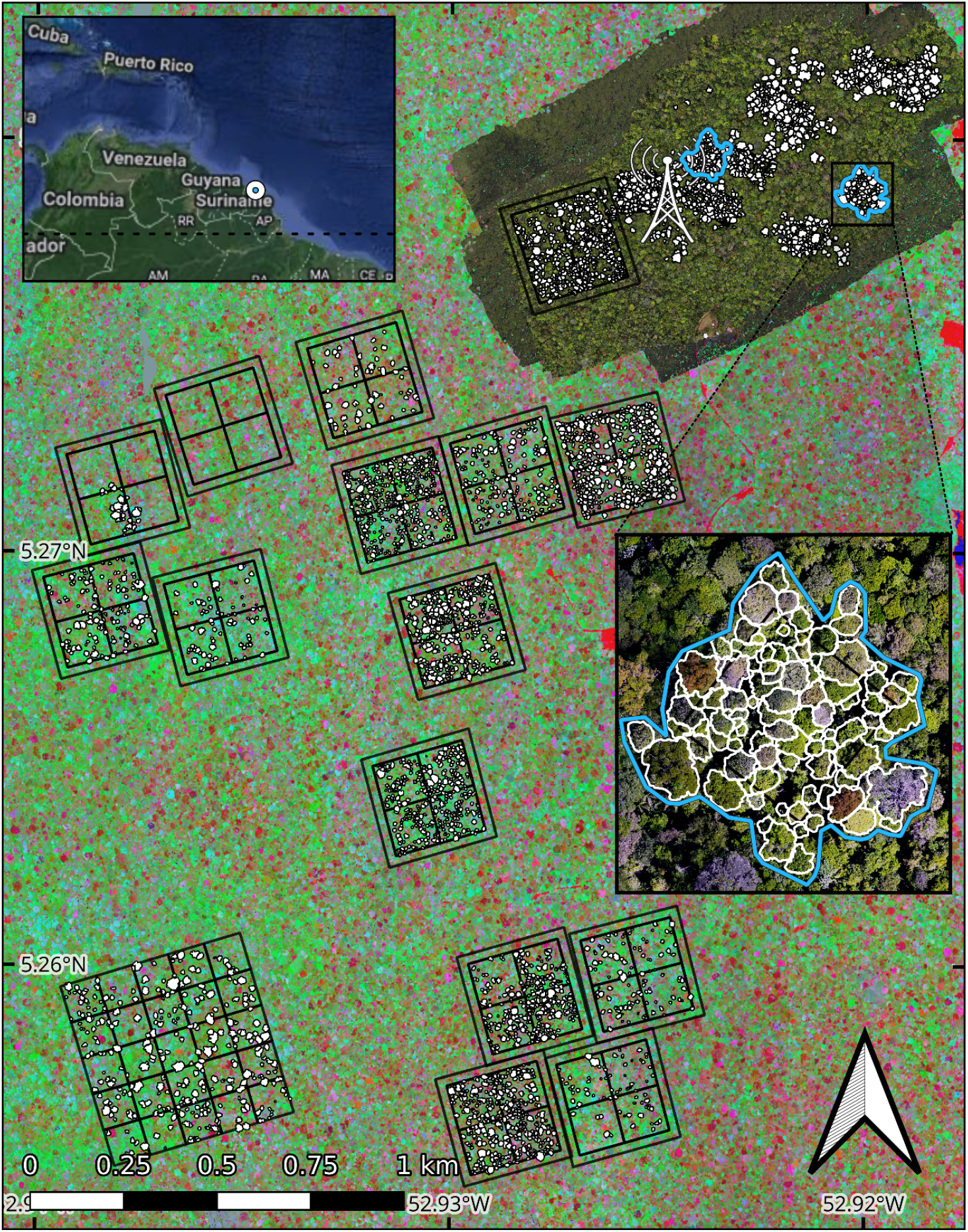
Study site at Paracou, French Guiana, showing the spatial distribution of manually delineated and labelled tree crowns (white outlines) overlaid on a hyperspectral composite (selected PCA bands). Permanent inventory plots are outlined in black. The UAV RGB survey region is shown in the northeast around the flux tower; the blue outlines are the crown-segmentation test areas used by Ball et al. [2026], one of which is enlarged at lower right. Adapted from that paper.

We used a reference database of 3,256 hand-delineated, field-verified tree crowns of 239 species, built over eight field missions between 2015 and 2023 [Ball et al., 2026]. Crowns were delineated in QGIS on the RGB, hyperspectral and LiDAR data and matched to inventory trees from the mapped trunk location and stem size [Gourlet-Fleury et al., 2004], with each match checked in the field against the imagery; every polygon carries confidence scores for crown integrity and for the trunk match, so geolocation uncertainty beneath the canopy is handled by field verification rather than by coordinate precision. Liana-infested crowns were retained so that infestation could be tested as a predictor; liana presence was recorded in the field when noticed rather than by systematic survey. Excluding the 70 species with a single crown left 3,186 crowns of 169 species, and pixels were filtered for illumination and crown footprint before classification (Methods S1).

### 2.2 Species classification from hyperspectral imagery

Airborne hyperspectral data were acquired on a single date, 19 September 2016, by two co-mounted HySpex sensors (VNIR-1600 and SWIR-384) covering 416–2500 nm at 1 m resolution (SWIR resampled from 2 m; Table 1). After removal of noisy SWIR bands, 378 of 448 channels were retained (preprocessing summarised in Methods S1; full details in Laybros et al. 2019 and Ball et al. 2026). Species were classified from pixel spectra (378 bands) by linear discriminant analysis (LDA), with each crown labelled from its pixels. LDA outperformed the more flexible classifiers tested on these data [Ball et al., 2026]. Because the hyperspectral data come from this single date, the main phenological metrics describe each species’ behaviour *across* the full annual cycle rather than its state on the acquisition date; whether that state also matters is tested separately as a follow-up.

**Table 1:** Remote sensing data sources (see Ball et al. 2026 for full acquisition and processing details). Resolution is ground resolution for RGB and processed canopy height model (CHM) resolution for LiDAR. The ten individual UAV RGB survey dates, and the aircraft used on each, are listed in Table S1. The 3.5-year UAV RGB phenology series (October 2020–May 2024, surveys about every three weeks) from which the leaf-cycle metrics derive is described with the phenology predictors in the Methods and in Ball [2023].

| Scan date(s) | Modality | Resolution | Altitude | Spectral range | Sensor |
| --- | --- | --- | --- | --- | --- |
| 23-Oct-2020–06-Apr-2021 | RGB | 5 cm | 70 m | 421–617 nm (3 bands) | DJI Phantom 4 Advanced; P4 Multispectral |
| 19-Sep-2016 | Hyperspectral | 1 m | 900 m | 416–992 nm (160 channels) | HySpex VNIR-1600 |
| 19-Sep-2016 | Hyperspectral | 2 m | 900 m | 930–2500 nm (288 channels) | HySpex SWIR-384 |
| 15-Nov-2019 | LiDAR | 0.5 m (CHM) | 800 m | 1064 nm (active) | RIEGL LMS-Q780 |

### 2.3 Sample-size standardisation and the 45-species dataset

Species had to be compared at equal sample size, so we first modelled per-species F1 as a function of the number of labelled crowns per species, using the per-species F1 of the full 169-species classification of Ball et al. [2026] (repeated stratified-group five-fold cross-validation over the 2,762 filtered crowns, with no crown split across folds; Methods S1), in which about four-fifths of a species’ crowns are used for training in each fold. We fitted a logistic learning curve with its lower asymptote at zero, F1(*n*) = *d/*(1 + exp(*−b* (log *n −* log *c*))), by nonlinear least squares, defined reliable performance as F1 ≥ 0.7, and computed the *n* at which the fitted curve crosses this threshold. Uncertainty was obtained by a bootstrap that resampled both species and cross-validation folds (2,000 resamples). On the basis of this curve, we included only species with at least ten crowns in the training partition (2,519 crowns of 169 species after pixel filtering), which left 49 species; removing four whose predictor data were incomplete left 45 species with 1,969 crowns.

All other analyses use new classification experiments with training data standardised to ten crowns per species. In each of 100 replicate analyses, we drew ten crowns per species at random and split them five times into eight training and two validation crowns per species (100 × 5 = 500 runs). An LDA was fitted in each run, with crowns labelled from the mean posterior probability of their pixels, and per-species F1 was averaged over the 500 runs. The validation-crown predictions give the confusion rates of the pairwise analysis and the outcomes of the crown-level analysis (Table 2).

**Table 2:**
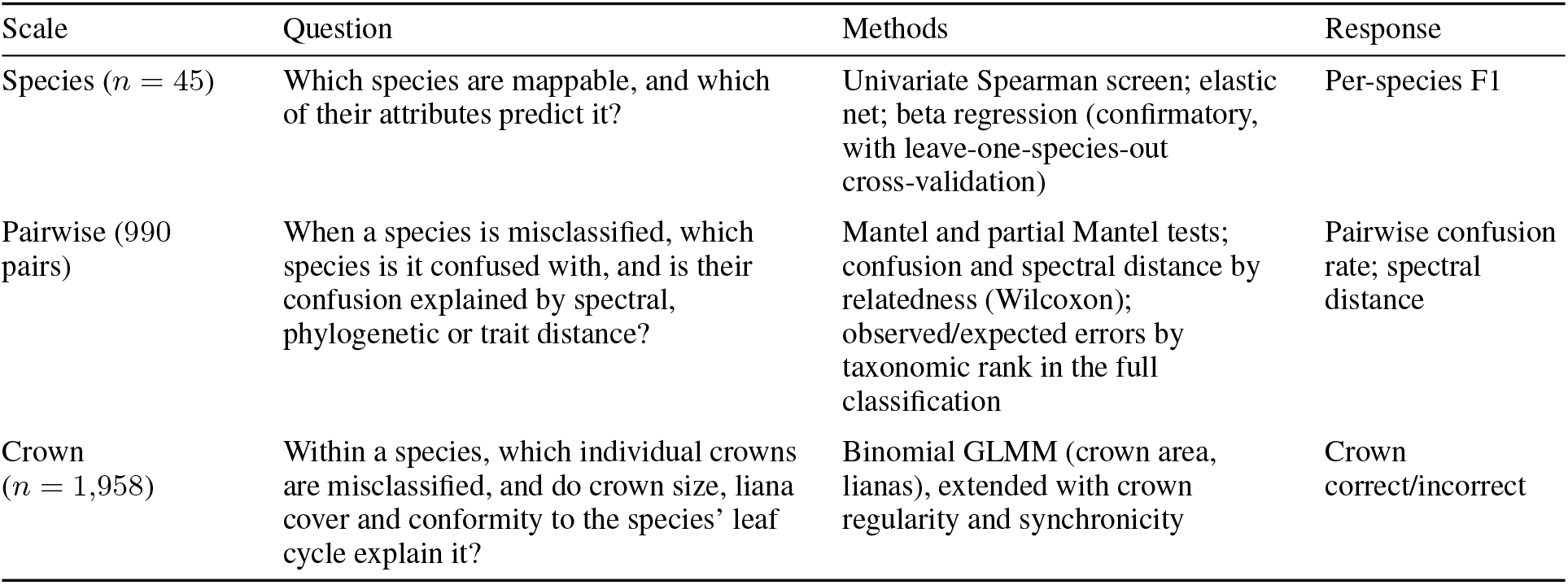
Three scales of analysis evaluating the drivers of species separability. The crown-level analysis uses the 1,958 of the 1,969 crowns of the 45 species that were drawn as validation crowns at least once (45,000 classifications <u>in total: 45 species</u> *<u>×</u>* <u>2 validation crowns</u> *<u>×</u>* <u>5 splits</u> *<u>×</u>* <u>100 replicates).</u>

| Scale | Question | Methods | Response |
| --- | --- | --- | --- |
| Species ($n = 45$) | Which species are mappable, and which of their attributes predict it? | Univariate Spearman screen; elastic net; beta regression (confirmatory, with leave-one-species-out cross-validation) | Per-species F1 |
| Pairwise (990 pairs) | When a species is misclassified, which species is it confused with, and is their confusion explained by spectral, phylogenetic or trait distance? | Mantel and partial Mantel tests; confusion and spectral distance by relatedness (Wilcoxon); observed/expected errors by taxonomic rank in the full classification | Pairwise confusion rate; spectral distance |
| Crown ($n = 1,958$) | Within a species, which individual crowns are misclassified, and do crown size, liana cover and conformity to the species' leaf cycle explain it? | Binomial GLMM (crown area, lianas), extended with crown regularity and synchronicity | Crown correct/incorrect |

### 2.4 Spectral distinctiveness

We described each species’ position in spectral space with three metrics computed from crown-level (median) spectra: *within-species spectral variability* (the standard deviation among conspecific crowns, averaged across the 378 standardised bands) and, from the Bhattacharyya distance [Bhattacharyya, 1943] between each pair of species, which combines the shift in mean spectra with differences in spectral dispersion, each species’ *spectral isolation* (mean distance to all other species) and *nearest-neighbour spectral distance* (distance to the most similar species), the spectral analogues of the phylogenetic metrics MPD and NNPD described below. Because covariance matrices cannot be estimated in 378 dimensions from tens of crowns, and distance estimates depend on sample size, all metrics were computed in the space of the first five principal components of the crown spectra (96.9% of variance), from ten randomly drawn crowns per species, averaged over 200 draws (Methods S1). Sensitivity analyses with Mahalanobis distances and with band-restricted metrics are described in Methods S1. They were analysed alongside, not among, the 29 candidate predictors described below.

### 2.5 Analytical framework for species separability

We quantified the influence of ecological, spatial and evolutionary factors on separability at three scales (Table 2) on the standardised 45-species dataset. The **species-level analysis** relates species-level F1 to the candidate predictors by a univariate Spearman screen, elastic-net variable selection and beta regression; the **pairwise analysis** asks whether the spectral distance between two species predicts how often they are confused (Mantel and partial Mantel tests); and the **crown-level analysis** models the classification outcome of each crown with a generalised linear mixed model (GLMM). A follow-up at the species level asked whether the timing of leaf-cycle events relative to the acquisition date adds anything to phenological regularity.

### 2.6 Candidate predictors

The 29 candidate predictors fall into the six groups of the Introduction.

**(1) Spatial distribution:** Spatial separation was measured by the mean pairwise distance from a 10-sample subset of individuals within a species, averaged over 1,000 random draws. Other clustering metrics were distorted by the non-uniform arrangement of the field plots and were not used.

**(2) Liana prevalence:** For each species, we calculated the proportion of crowns recorded as having lianas in the field surveys.

**(3) Crown size and tree height:** Crown area (polygon area) and tree height (median CHM value within the crown) were tested as predictors.

**(4) Leaf and bark traits:** We used the 12 species-mean traits available for all 45 species in Vleminckx et al. [2021]: leaf nitrogen, carbon, *δ*^13^C, phosphorus, calcium, potassium, chlorophyll content, toughness, area, thickness, specific leaf area (SLA), and trunk bark thickness. All 12 entered the univariate screen and elastic net individually. Because spectral and phylogenetic predictors are expressed as distinctiveness, we also computed two distinctiveness metrics in the z-scored 12-trait space by direct analogy with NNPD and MPD: nearest-neighbour trait distance and trait isolation (mean Euclidean distance to all other species). Each was tested by adding it to the confirmatory beta regression and by substituting the pair for the 12 traits in the elastic net; leaf carbon, the trait the elastic net weighted most heavily, was tested in the same way as an exploratory check.

**(5) Leaf phenology:** We used six metrics derived from a 3.5-year time series (October 2020–May 2024) of crown-level greenness (Green Chromatic Coordinate, GCC) measured every three weeks from repeat UAV surveys and interpolated to daily values (Ball 2023; illustrated in Fig. S1). Three describe how closely conspecific crowns share a leaf cycle: (i) *synchronicity*, the mean lag-zero correlation between the greenness series of conspecific crowns, which is high when the trees of a species are in step; (ii) *regularity*, the maximum cross-correlation between the same series over lags of up to ±365 days, averaged over all conspecific pairs, which is high when conspecifics share a greenness waveform whether or not they are in step; and (iii) *lag*, the mean absolute lag (days) at which that maximum occurs. Three further descriptors of the greenness cycle, amplitude, period and seasonality concentration (Methods S3), were included as candidate predictors. Regularity represents phenology in the beta regression because it predicted F1 better than synchronicity or any composite of the six metrics.

**(6) Phylogenetic relatedness:** We computed seven phylogenetic metrics on the time-calibrated phylogeny: nearestneighbour patristic distance (NNPD), nearest-congener distance (zero for species without a congener), terminal branch length, evolutionary distinctiveness (fair proportion and equal splits), mean patristic distance (MPD), and a binary indicator for the absence of congeners among the species of the crown dataset. NNPD captures proximity to the closest relative; MPD summarises overall phylogenetic isolation (Methods S2).

### 2.7 Species-level (one-vs-all separability) analyses

The species-level analysis proceeds in three steps on the 45-species standardised dataset. First, a univariate screen: Spearman correlations of each of the 29 candidate predictors with F1, with partial correlations controlling for crown size, greenness amplitude or the number of crowns per species where an association could reflect them (ppcor). Second, variable selection by elastic net: penalised regression of logit-transformed F1 on all 29 predictors (glmnet; Friedman et al. 2010), with ten-fold cross-validation choosing the mixing parameter *α* and the regularisation parameter *λ*. Third, confirmatory beta regression [Ferrari and Cribari-Neto, 2004] with a logit link on Smithson–Verkuilen-transformed [Smithson and Verkuilen, 2006] F1, using a small set of standardised predictors specified a priori, at most one per factor group: M1 (spatial dispersion), M2 (+ regularity), M3 (+ liana prevalence), and M4–M7, each adding one term to M3 (MPD, tree height, within-species spectral variability, spectral isolation); M6, M7 and a model with both spectral terms combine spectral and ecological predictors. Models were compared by AIC and by leave-onespecies-out (LOSO) cross-validation, and coefficients were tested by permuting the response (1,000 permutations). The three are parallel analyses of the same data rather than a pipeline: they use different predictor sets, so they are compared by whether they agree on which factors matter, not by coefficient values.

### 2.8 Pairwise separability analyses

Pairwise confusion was the mean of the two directed crown-level confusion rates between each pair of species in the standardised classification. We tested (i) whether spectral distance (Bhattacharyya) predicted confusion, using a Mantel test (Spearman’s *ρ*, 9,999 permutations, two-sided); (ii) whether this relationship persisted after controlling for the patristic distance between the two species (partial Mantel test); and (iii) whether distances between species in phenology (six metrics), traits (12 traits), phylogeny (patristic distance) and stature (height, crown area) predicted spectral distance or confusion. Phenology, trait and stature distances were Euclidean distances on z-scored variables. We also compared confusion and spectral distance among congeneric pairs, other confamilial pairs, and pairs from different families (Wilcoxon rank-sum tests). Finally, for the predictions on the held-out test set of 666 crowns of the full 169-species classification of Ball et al. [2026], we counted the misclassified crowns assigned to a congener, to another species of the same family, or to a species of a different family, and compared each count with its expectation under random reassignment among the other species in the pool (Methods S2). Mantel tests used vegan [Oksanen et al., 2022].

### 2.9 Crown-level analyses

To identify crown attributes that influence correct classification, we used the crown-level predictions from the standardised classification (45,000 classifications of 1,958 crowns; Table 2). Each crown was classified repeatedly (median 14 times), and its outcomes were strongly correlated: 65% of crowns were always classified correctly and 9% never. We therefore analysed crowns rather than individual classifications, taking as the response whether a crown was classified correctly in the majority of its classifications. We fitted a binomial GLMM with crown area (log-transformed, standardised) and liana presence (binary) as fixed effects and a random intercept for species:

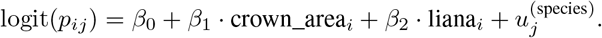

The model was fitted using lme4::glmer [Bates et al., 2015], and marginal and conditional *R*^2^ were computed following Nakagawa and Schielzeth [2013]. Species random intercepts were correlated with species-level F1 to check that the two scales agree. A quasi-binomial model of the proportion of each crown’s classifications that were correct gave the same conclusions.

To ask whether phenology also acts at the level of the individual crown, we computed each crown’s own regularity and synchronicity: the mean, over its conspecifics in the UAV series, of the maximum (regularity) or lag-zero (synchronicity) cross-correlation between the two greenness series, so that the species-level metrics of (5) are the means of these crown values. For the 535 crowns present in both the standardised classification and the phenology series, we added crown regularity or synchronicity, centred within species and standardised, to the GLMM above, together with the crown integrity score from the field database as a check on whether phenological consistency merely indexes crown quality (Methods S3).

### 2.10 Follow-up at the species level: leaf-cycle timing and single-date homogeneity

We tested whether the phenological state of a species on the acquisition date, or its uniformity among conspecifics, predicted separability. From the circular mean flush and shed dates of each crown, and their species means, we computed leaf age at scan, its standard deviation among conspecific crowns, and the proximity of flush and shed to the acquisition date, and, as a measure that does not depend on identifying flush events, the standard deviation of crown-standardised greenness among conspecific crowns on the acquisition calendar date (definitions in Methods S3). Each metric was tested by Spearman correlation with F1, and timing metrics were also substituted for, or added to, regularity in model M3.

## 3 Results

### 3.1 The learning curve and the standardised dataset

The fitted learning curve of F1 against the number of labelled crowns reached an expected F1 of 0.7 at 17 crowns per species (bootstrap 95% CI [14, 21]) and levelled off at F1 *≈* 0.81 (Fig. 2). Few species with few crowns were classified reliably: 9 of the 136 species with fewer than 20 crowns reached F1 ≥ 0.7, against 29 of the 33 species with 20 or more. Training data are strongly skewed towards a few common species (Fig. S2): 135 of 169 species (80%) have fewer than 17 crowns. Because expected F1 is still rising steeply across this range, the comparisons that follow hold sample size constant. Ten crowns per species is a compromise that serves both ends: it lies on the steep part of the curve, below saturation, so that differences among species are not flattened by abundant training data, and it is reached by enough species (49) to support the analyses.

**Figure 2:**
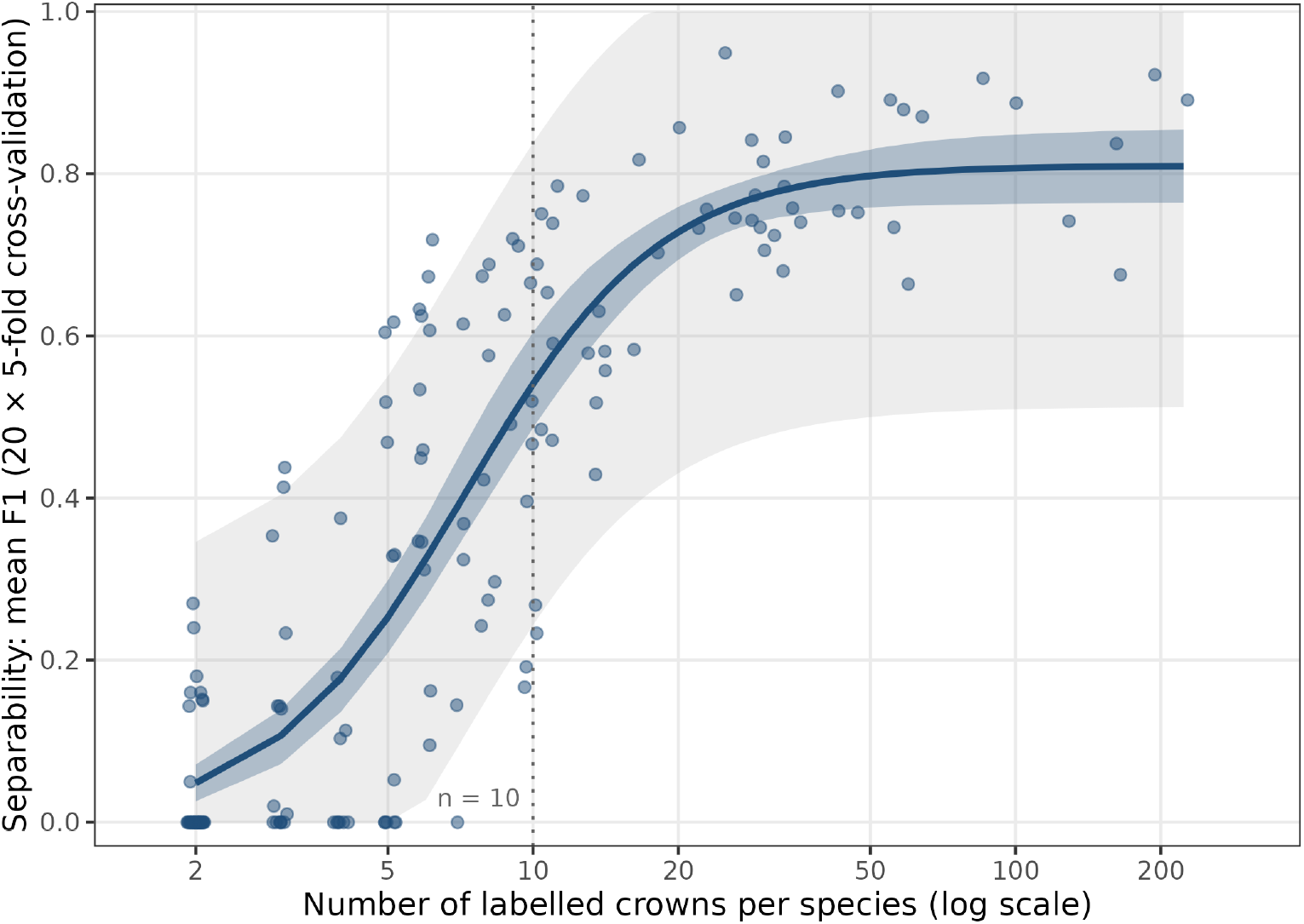
The learning curve: mean F1 against the number of labelled crowns per species (20 × 5-fold cross-validation, all 169 species, log-*x* axis, points jittered). Logistic fit with its lower asymptote at zero; inner band, bootstrap 95% CI; outer band, ±1.96 residual SD. The dotted line marks *n* = 10, the sample size of all later analyses. Per-species ranking and confusion matrix in Fig. S4.

With sample size equalised across the 45 species of the standardised dataset, F1 still ranged from 0.60 (*Licania alba*) to 0.99 (*Manilkara bidentata*), with 38 of the 45 species above 0.7 (Fig. S3), so the differences among species that the determinants must explain are large.

### 3.2 Spectral distinctiveness

As expected, separability tracked spectral distinctiveness closely, a result that is as much a check on the classifier as a finding. In the standardised dataset (per-species F1 in Fig. S3), species that were spectrally distant from other species and internally consistent were easier to classify (Fig. 3; Fig. S5; examples in Figs. S6 and S7). F1 increased with spectral isolation (*ρ* = 0.45, *p* = 0.002) and with nearest-neighbour spectral distance (*ρ* = 0.38, *p* = 0.010), and decreased with within-species spectral variability (*ρ* = −0.31, *p* = 0.038). The two faces of distinctiveness are visible in Fig. 3a,b. The three most separable species, *Manilkara bidentata*, *Xylopia nitida* and *Couma guianensis* (F1 = 0.99), each form a tight cluster of crowns, whereas the three least separable, *Licania alba*, *Tapura capitulifera* and *Inga alba* (F1 0.60–0.64), spread across the whole cloud of crowns, so that no region of spectral space is characteristic of them. A species is separable, in other words, when its crowns resemble one another more than they resemble those of other species, and the gradient between these extremes is continuous (Fig. 3c). Band by band, variability among conspecific crowns was negatively related to F1 throughout the spectrum, most strongly at the red edge (*ρ* = −0.53 at 709 nm; Fig. S8).

**Figure 3:**
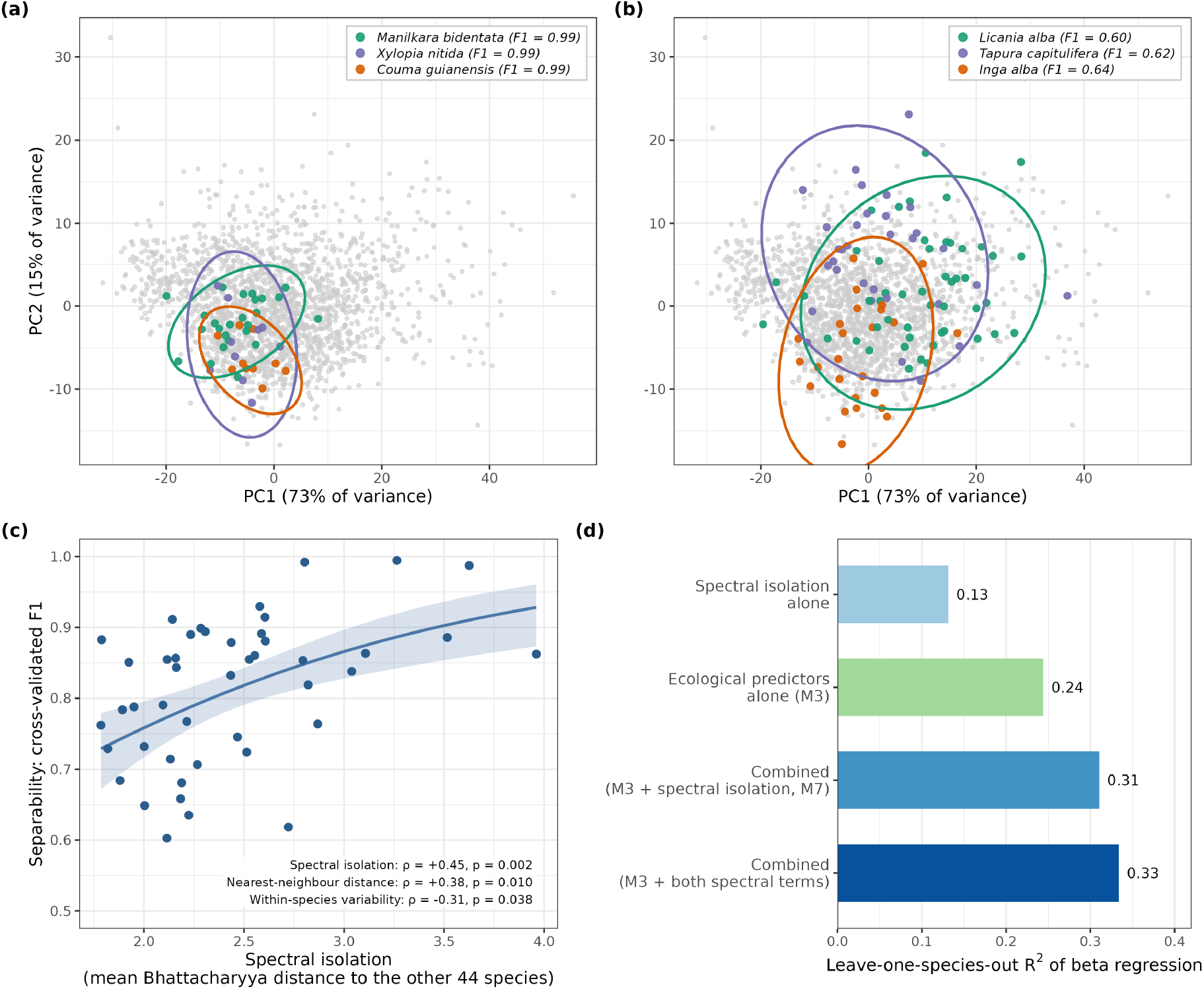
Spectral distinctiveness (*n* = 45 species). (a, b) Crown spectra in the space of their first two principal components (73% and 15% of variance); grey points are all crowns, and the three most (a) and least (b) separable species are highlighted with 95% concentration ellipses. (c) Species-level F1 against spectral isolation (smooth fit with 95% band) with the Spearman correlations of the three spectral metrics. (d) Leave-one-species-out *R*^2^ of beta regressions with spectral isolation alone, the ecological model M3 (dispersion, regularity, lianas), both combined (M7) and M3 with both spectral terms; full model set in Fig. S9.

Restricting the spectra to the 10% or 25% of bands most important for classification, under either ranking, did not strengthen the relationships of F1 with spectral isolation or nearest-neighbour distance, and changed the other correlations by at most 0.09 (Table S2): with the top 38 bands by LDA scalings, which include all eight far-red-edge (748–775 nm) bands, the correlation of F1 with spectral isolation was 0.41 (0.45 with all bands) and the Mantel correlation between spectral distance and confusion −0.30 (−0.32). A tenth of the bands, chosen by either criterion, carries nearly as much species-level information as the full spectrum, and no more.

### 3.3 Species-level determinants of separability

We report the six determinants in the order of the Introduction, giving for each the univariate Spearman correlation with F1 (all 29 predictors in Fig. S10), whether the elastic net retained it, and its coefficient in the confirmatory beta regression (Fig. 4). The three methods agreed on the main ecological and evolutionary predictors. The elastic net retained 8 of 29 predictors: spatial dispersion, liana prevalence, regularity, seasonality concentration, absence of congeners, MPD, leaf carbon and bark thickness, in the order of Fig. 4, with absence of congeners, MPD and spatial dispersion the largest. The confirmatory model M3 (spatial dispersion + regularity + liana prevalence) had pseudo-*R*^2^ = 0.29 and LOSO *R*^2^ = 0.24 (Figs. S9 and S11); adding MPD (M4) raised these to 0.38 and 0.27, and M4, whose partial effects are shown in Fig. 5, is the confirmatory model shown in Fig. 4. Spectral distinctiveness predicted F1 more strongly still and was largely complementary to them (Fig. 3d; last paragraph of this section).

**Figure 4:**
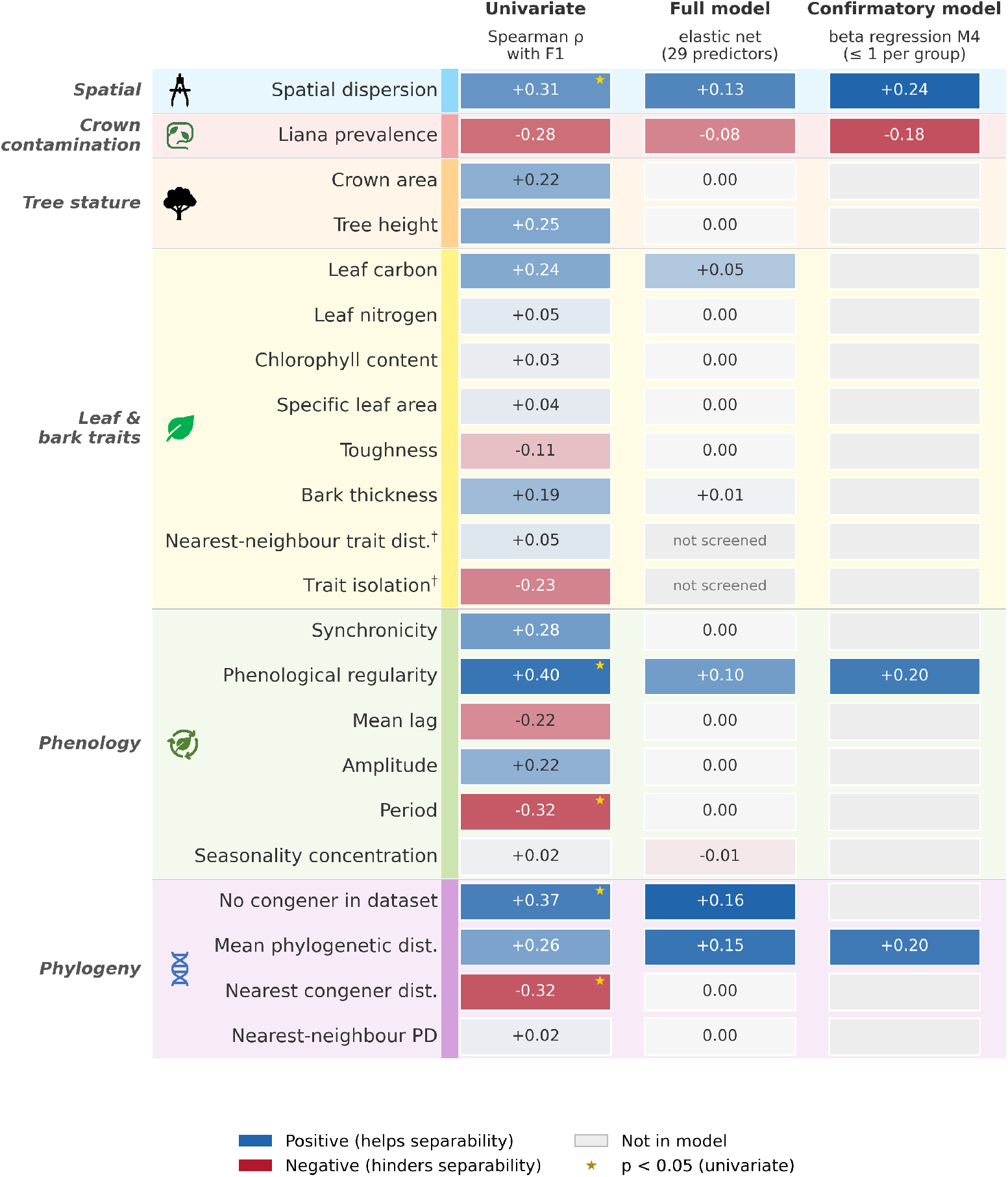
Species attributes as predictors of separability (*n* = 45 species). Rows are the 20 predictors with a non-zero result in any analysis, grouped as in the Introduction (symbols); the nine omitted (three phylogenetic metrics, six traits) had zero coefficients throughout. Columns: univariate Spearman correlation with F1 (stars, *p <* 0.05); elastic net over all 29 predictors (0.00 = dropped by the penalty); confirmatory beta regression M4, specified a priori with one representative per group (dispersion, regularity, lianas, MPD). Colour gives the sign and intensity the magnitude relative to the largest value in the column; grey tiles are predictors not in that model. *†* Trait-distinctiveness metrics added post hoc, not screened by the elastic net (Table S3).

**Figure 5:**
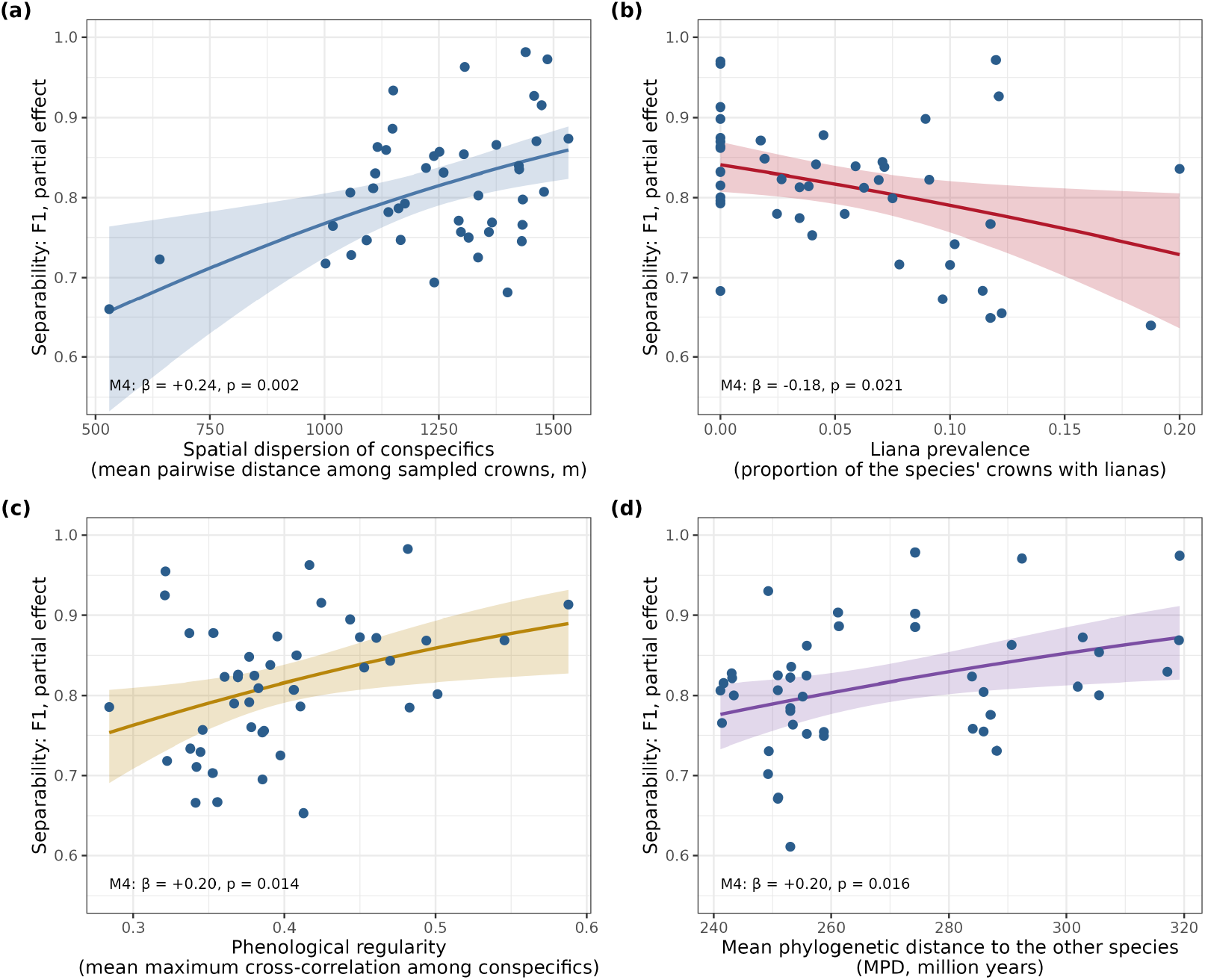
Partial effects of the four predictors of the confirmatory model M4 on species-level F1 (*n* = 45 species): (a) spatial dispersion, (b) liana prevalence, (c) phenological regularity, (d) mean phylogenetic distance. Lines are M4 predictions with the other three predictors at their means (95% confidence bands); points are partial residuals, each species’ F1 adjusted for the other three predictors; *β* is the standardised coefficient with its Wald *p*. Univariate correlations and elastic-net coefficients of the same predictors are in Fig. 4.

**(1) Spatial distribution:** Species whose individuals were more widely dispersed across the landscape were classified more accurately, and the three methods agreed: *ρ* = 0.31 (*p* = 0.041), one of the three largest elastic-net coefficients, and the largest and most stable coefficient in the beta regressions (+0.28 in M3, *z* = 3.56, permutation *p* = 0.007; +0.24 in M4), with or without spectral terms.

**(2) Liana presence:** Species with higher rates of liana infestation were harder to classify. The univariate correlation was marginal (*ρ* = −0.28, *p* = 0.063), but the elastic net retained liana prevalence and it was significant in the beta regression (−0.21 in M3, *z* = −2.63, permutation *p* = 0.038; −0.18 in M4); adding it raised LOSO *R*^2^ from 0.18 in M2 to 0.24 in M3. The effect was confirmed at the crown level.

**(3) Tree stature and crown size:** No method supported stature at the species level. Neither mean tree height (*ρ* = 0.25, *p* = 0.10) nor mean crown area (*ρ* = 0.22, *p* = 0.14) was correlated with F1, the elastic net dropped both, and adding tree height to M3 (M5) did not improve prediction (LOSO *R*^2^ = 0.20). At the crown level, however, larger crowns were classified more reliably.

**(4) Leaf and bark traits:** No trait was significantly correlated with F1 (Table S4; Figs. S12 and S13); the strongest association was with leaf carbon (*ρ* = 0.24, *p* = 0.11). Recasting traits as distinctiveness did not change this: neither nearest-neighbour trait distance (*ρ* = 0.05, *p* = 0.73) nor trait isolation (*ρ* = −0.23, *p* = 0.12; the sign opposite to that expected if distinctive traits aided separability) was correlated with F1, adding either to M3 lowered LOSO *R*^2^ (0.24 to 0.22; permutation *p* = 0.79 and 0.55), substituting the pair for the 12 traits in the elastic net left the other selections unchanged (Table S3), and distance in trait space between two species did not predict how often they were confused. The elastic net retained leaf carbon and bark thickness with small coefficients, but adding leaf carbon to M3 reduced LOSO *R*^2^ (0.24 to 0.17), as expected for a predictor selected from 12 correlated candidates using the response. The traits as a group add nothing that generalises: the 12 traits together had an in-sample adjusted *R*^2^ of 0.23 but a LOSO *R*^2^ of −0.20, and 12 random predictors reach the same adjusted *R*^2^ in 5% of simulations.

**(5) Phenological regularity:** Because classification used a single acquisition date, synchronicity, which requires conspecifics to be in phase, might be expected to matter more than regularity, which does not. It was regularity, however, on which the methods agreed (Fig. 5c). Species whose crowns shared a pronounced, consistent greenness cycle were easier to classify: regularity had the strongest univariate correlation of the 29 predictors (*ρ* = 0.40, *p* = 0.006), was retained by the elastic net (fourth by coefficient), and was retained in M3 (+0.21, *z* = 2.37, permutation *p* = 0.050) and M4 (+0.20). The association was robust to controlling for crown area and tree height (partial *ρ* = 0.37, *p* = 0.014), greenness amplitude (partial *ρ* = 0.39, *p* = 0.009), or the number of crowns per species (partial *ρ* = 0.43, *p* = 0.003), and it was mirrored within species at the level of the individual crown.

The two metrics were strongly correlated (*ρ* = 0.72, *p <* 0.001; Fig. S14), as expected since synchronicity is a lower bound on regularity, even though lags between conspecific crowns were substantial (mean 120 days across species, range 39–202; Fig. S15). Regularity was unrelated to the number of crowns a species contributed (*ρ* = 0.13, *p* = 0.39) and was higher in species whose conspecific crowns had shorter lags (*ρ* = −0.50, *p <* 0.001); lag itself was a weak predictor of F1 (*ρ* = −0.22, *p* = 0.16; Fig. 4). Of the remaining metrics, only period was related to F1 (*ρ* = −0.32, *p* = 0.031; Fig. 4). Regularity nonetheless carried the association with F1: it remained associated with F1 after controlling for synchronicity (partial *ρ* = 0.30, *p* = 0.046), whereas synchronicity did not after controlling for regularity (partial *ρ* = −0.02, *p* = 0.92), and substituting synchronicity for regularity in M3 reduced LOSO *R*^2^ from 0.24 to 0.17. The evidence is modest, however: the difference between the two univariate correlations (0.40 and 0.28) was not itself significant (Williams’ test, *p* = 0.25), and the two metrics were equally reliable as species-level estimates (split-half reliability 0.66 and 0.69; Methods S3), so the advantage of regularity is not explained by lower measurement noise.

**Single-date homogeneity and leaf-cycle timing:** Measures of how uniform a species was on the acquisition date did not predict F1. The spread of greenness among conspecific crowns on the acquisition calendar date tracked synchronicity (*ρ* = −0.44, *p* = 0.002) but not F1 (*ρ* = −0.14, *p* = 0.36), and neither leaf age at scan (*ρ* = −0.12; Fig. S16), leaf-age SD (*ρ* = −0.01; Fig. S17), flush proximity (*ρ* = −0.07) nor shed proximity (*ρ* = −0.11; all *p >* 0.4; flush and shed dates relative to the acquisition in Fig. S18) was related to F1. Substituting any timing metric for regularity in M3 gave a poorer model (LOSO *R*^2^ ≤ 0.19), and adding them to regularity did not improve it. Regularity was also unrelated to within-species spectral variability (*ρ* = 0.08, *p* = 0.59; Fig. S19). Regular species were therefore not more uniform on the day of acquisition; they were, on average, further from other species in spectral space.

**(6) Phylogenetic relatedness:** The 25 species with a congener in the crown dataset were harder to classify than the 20 without (mean F1 0.78 vs. 0.86; *ρ* = 0.37, *p* = 0.012 for congener absence), and the contrast was at least as strong when only congeners classified in the same pool were counted (16 species with such a congener; *ρ* = −0.39, *p* = 0.008 for congener presence). Among species with congeners, the phylogenetic distance to the nearest one was unrelated to F1 (*ρ* = 0.03, *p* = 0.89), so it is the presence of a congener, not its closeness, that matters. Species without congeners were also more spectrally isolated (*ρ* = 0.34, *p* = 0.023). MPD was not significant alone (*ρ* = 0.26, *p* = 0.085) but was retained by the elastic net, with one of its three largest coefficients, and improved the beta regression (M4, +0.20). F1 itself showed no phylogenetic signal (Pagel’s *λ* = 0.24, *p* = 0.28; Blomberg’s *K* = 0.013, *p* = 0.52).

**Combining spectral and ecological predictors:** Adding spectral terms to M3 improved prediction (Fig. 3c,d). LOSO *R*^2^ rose to 0.26 with within-species spectral variability (M6), to 0.31 with spectral isolation (M7), and to 0.33 with both (pseudo-*R*^2^ = 0.50), whereas spectral isolation alone gave LOSO *R*^2^ = 0.13. Spectral and ecological predictors were therefore largely complementary. In the model with both spectral terms, spatial dispersion kept its effect (*β*^^^ = +0.25, permutation *p* = 0.014), while the coefficients of regularity (+0.19, *p* = 0.092) and liana prevalence (−0.14, *p* = 0.24) were smaller than in M3. Regularity was positively correlated with spectral isolation (*ρ* = 0.29, *p* = 0.057) and with nearest-neighbour spectral distance (*ρ* = 0.31, *p* = 0.037), so part of its association with F1 is shared with spectral distinctiveness.

### 3.4 Pairwise analysis: spectral distance and confusion

Pairs of species that were closer in spectral space were confused more often (Mantel *r* = −0.32, *p <* 0.001; Fig. 6a), and this relationship was unchanged after controlling for the patristic distance between them (partial Mantel *r* = −0.32, *p <* 0.001). Distances between species in phenology, traits or stature predicted neither spectral distance nor confusion (Table 3). Phylogenetic distance predicted confusion (*r* = −0.19, *p* = 0.002), also after controlling for spectral distance (partial *r* = −0.19, *p* = 0.002), although across all pairs it did not predict spectral distance.

**Figure 6:**
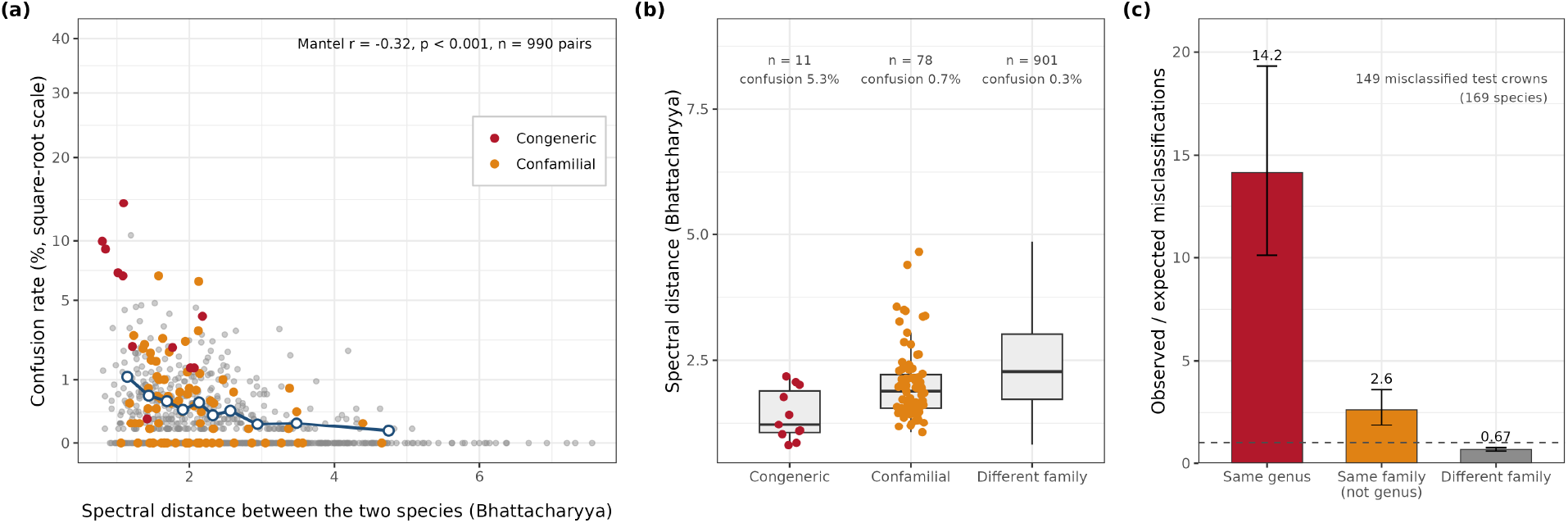
Pairwise separability among the 45 standardised species (990 pairs). (a) Confusion rate against spectral (Bhattacharyya) distance (square-root axis; Mantel *r* on the panel); congeneric and confamilial pairs coloured, blue points the mean confusion per decile of distance. (b) Spectral distance by relatedness of the pair (median and interquartile range; coloured points are the congeneric and confamilial pairs), with the number of pairs and their mean confusion above each box. (c) Observed/expected misclassifications by relatedness for the 149 misclassified test crowns of the full 169-species classification, against random assignment among the other 168 species; Wilson 95% CIs; dashed line, null ratio of 1.

**Table 3:** Mantel tests among the 45 species of the standardised dataset (990 pairs; Mantel *r* on Spearman ranks, 9,999 permutations, two-sided *p*): do distances between species in each predictor group predict their spectral (Bhattacharyya) distance, or the rate at which they are confused? For comparison, spectral distance itself predicts confusion with *r* = −0.32 (*p <* 0.001).

| Predictor distance | Spectral distance |  | Confusion rate |  |
| --- | --- | --- | --- | --- |
| | Mantel $r$ | $p$ | Mantel $r$ | $p$ |
| Phenology (6 metrics) | +0.13 | 0.078 | -0.03 | 0.62 |
| Leaf and bark traits (12 traits) | +0.11 | 0.13 | -0.08 | 0.24 |
| Phylogeny (patristic distance) | +0.05 | 0.49 | -0.19 | 0.002 |
| Stature (height, crown area) | +0.06 | 0.42 | +0.01 | 0.87 |

The phylogenetic effect was confined to close relatives. Confusion averaged 5.3% for the 11 congeneric pairs, 0.7% for the 78 other confamilial pairs, and 0.3% for the 901 pairs from different families, and mean spectral distance rose in the same order (1.4, 2.0 and 2.5; Fig. 6b; both contrasts with between-family pairs Wilcoxon *p <* 0.001). Among pairs from different families, patristic distance was unrelated to confusion (*r* = −0.11, *p* = 0.12). The same structure appears in the errors of the full 169-species classification (Fig. 6c; Fig. S20): of the 149 misclassified test crowns, 29 were assigned to a congener, 14 times more than expected if errors fell at random among the other species (95% CI [10, 19]), and 28 to another species of the same family, 2.6 times more than expected (95% CI [1.9, 3.6]).

### 3.5 Crown-level analysis

Two crown attributes affected whether a crown was classified correctly (Fig. 7). Larger crowns were more likely to be identified correctly (*β*^^^ = +0.38 per SD of log crown area, *z* = 5.3, *p <* 0.001; odds ratio 1.44 per doubling of crown area; Fig. 7a): 76% of crowns in the smallest quartile (median 31 m^2^) were classified correctly, against 87–90% in the two largest quartiles. Liana presence more than halved the odds of correct classification (*β*^^^ = −0.82, *z* = −3.5, *p <* 0.001; odds ratio 0.44 [0.28, 0.70]; Fig. 7b): 72% of the 113 liana-infested crowns were classified correctly, against 85% of the others. Neither effect was significant at the species level.

**Figure 7:**
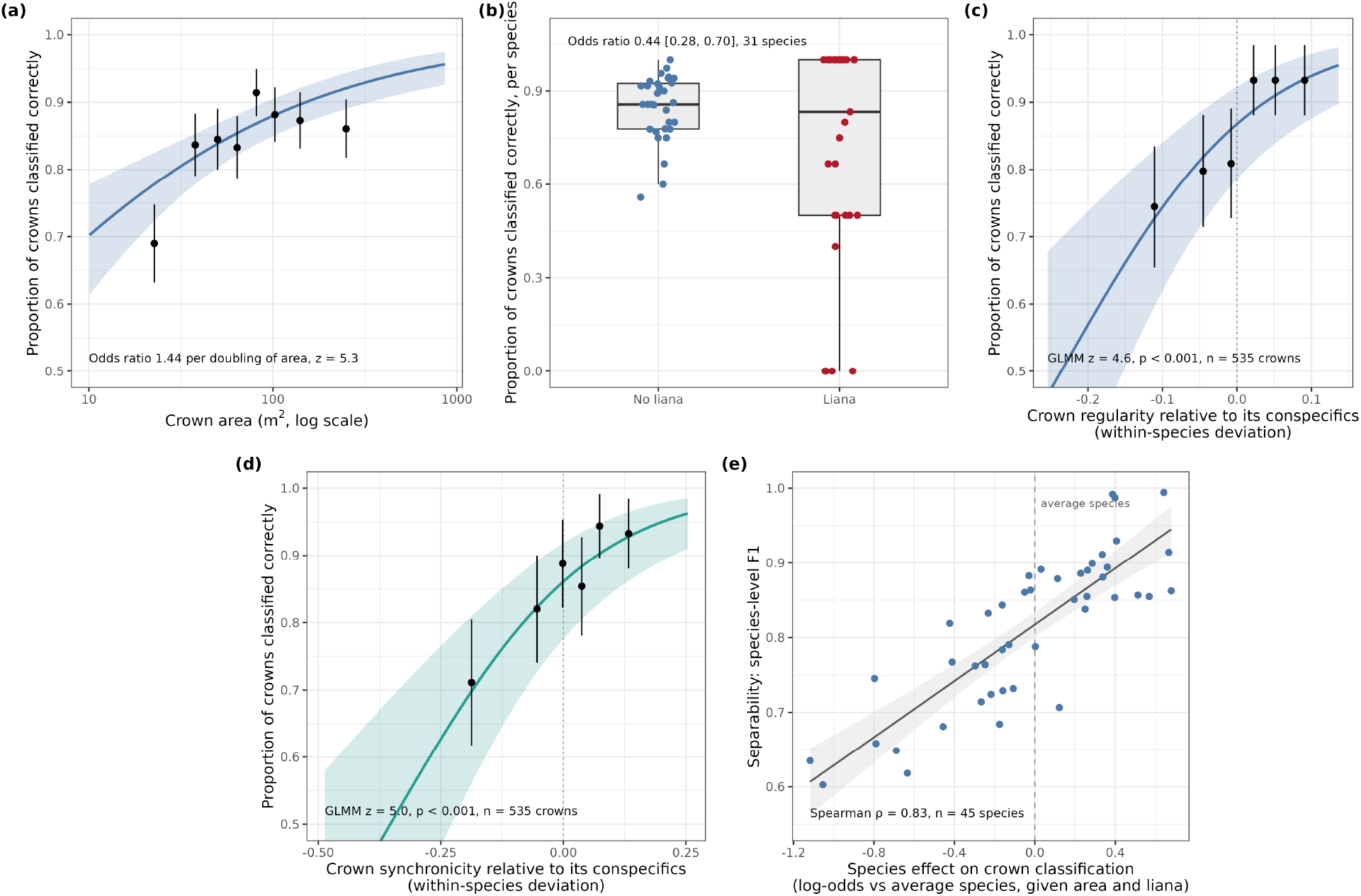
Crown-level drivers of correct classification (a crown is correct if correct in the majority of its classifications). (a, b) Binomial GLMM on 1,958 crowns of 45 species. (a) Proportion correct per octile of crown area (points, 95% CI) and the GLMM prediction for liana-free crowns (line, 95% band). (b) Proportion correct per species for crowns without and with lianas, in the 31 species with both (points jittered; boxes, median and interquartile range); the odds ratio is the GLMM estimate adjusted for crown area and species. (c, d) Whether a crown was classified correctly against its own (c) regularity and (d) synchronicity relative to its conspecifics, for the 535 crowns with a greenness series (points, proportion correct per sextile, 95% CI; line, GLMM prediction for a liana-free crown of average area, 95% band; *z*, the within-species term). (e) Each species’ random intercept in the crown-level GLMM (log-odds relative to the average species; dashed line at zero) against its species-level F1, with a linear fit and 95% band.

The marginal *R*^2^ of the model was 0.05 and the conditional *R*^2^ 0.14 (species random-effect SD = 0.61). Species random intercepts were strongly correlated with species-level F1 (Spearman *ρ* = 0.83, *p <* 0.001; Fig. 7e), showing that the two scales of analysis agree.

Phenological regularity also operated at the level of the individual crown. Among the 535 crowns with both a classification outcome and a greenness series (86% classified correctly), a crown’s own regularity relative to its conspecifics predicted whether it was classified correctly (*β*^^^ = +0.64 per within-species SD, *z* = 4.6, *p <* 0.001; AIC 437 to 416; Fig. 7c), whereas in this subset crown area, liana presence and the integrity score did not (*|z| ≤* 1.1). Crown synchronicity (*r* = 0.72 with crown regularity within species) predicted at least as well (*z* = 5.0; AIC 412; Fig. 7d) and retained the larger coefficient when both were included (*z* = 2.7 vs. 1.8); species-level regularity added nothing to a model containing crown regularity (*z* = 1.1). Phenological consistency was only modestly related to the two available indicators of crown quality: within species, lower-integrity crowns (−0.24 within-species SD, *t* = −3.0) and liana-infested crowns (−0.31 SD, *t* = −2.3) were somewhat less regular; across species, the share of lower-integrity crowns was unrelated to regularity (*ρ* = −0.22, *p* = 0.14), and the association of F1 with regularity was unchanged after controlling for it, liana prevalence and crown area (partial *ρ* = 0.44, *p* = 0.003).

## 4 Discussion

### 4.1 Signal consistency, not any single trait, governs mappability

Our three scales of analysis show that classification accuracy in this hyperdiverse forest is governed by the consistency and quality of the spectral signal a species presents rather than by the identity of any single functional trait (Fig. 4). Whether a species can be mapped depends first on how many labelled crowns are available and, once that is accounted for, on how far it sits from other species in spectral space and how consistently its crowns reproduce that spectrum. Among the species attributes, phenological regularity, spatial dispersion of training individuals and freedom from lianas were retained by every multivariate analysis, congeners in the classification pool impaired accuracy, and leaf chemical and structural traits, whether expressed as values or as distinctiveness, had no practical predictive value.

Spectral distinctiveness is the proximate quantity through which every attribute must act, not a rival explanation, and its correlation with F1 is in part a check that the classifier behaves as expected. The informative result is that the ecological attributes explained variation in F1 that spectral isolation alone did not, and that the two together predicted F1 out of sample better than either (Fig. 3d), so the attributes are not proxies for where a species sits in spectral space. Ball et al. [2026] found that the classifier gave most weight to the far-red edge (748–775 nm), a region sensitive to leaf ontogeny, stress and water content [Thomas and Gausman, 1977, Gitelson et al., 2003, Clark and Roberts, 2012], which is broadly consistent with, though not a test of, the role of leaf demography suggested below, and one in which reflectance carried phylogenetic signal here (Fig. S21; Methods S5; cf. Cavender-Bares et al. 2016). Yet specieslevel information was not concentrated there: metrics computed from the 38 bands weighted most heavily by the classifier, from a largely different set of bands in the shortwave infrared, or from the full spectrum predicted F1 and confusion about equally well. Reflectance is strongly correlated across bands, so the information that separates species is repeated across spectral regions rather than diluted by them, and band importance in a classifier should not be read as the location of the biological signal.

### 4.2 Why do regularity and synchronicity predict species separability?

Classification accuracy was expected to depend on phenological synchronicity, since conspecifics at the same developmental stage on the date of imaging should look alike. Our analyses show instead that regularity, fidelity to a shared leaf cycle whatever its timing, was the stronger predictor: neither synchronicity nor any single-date measure of uniformity predicted F1 once regularity was accounted for, and regular species varied among themselves as much as irregular ones but lay further from other species in spectral space. We hypothesise that regularity reflects leaf demography: it is highest in species whose crowns turn over their leaves in cohorts and lowest under continuous leaf exchange. It is distinct from deciduousness, being unrelated to field-observed leaflessness, which the amplitude of the greenness cycle tracks instead (Methods S3). Cohort renewal would enhance distinctiveness in two ways: a crown renewed in cohorts holds foliage of a narrow age range and so expresses the optical properties of its species at one leaf stage [Chavana-Bryant et al., 2017], whereas continuous turnover averages age classes and pulls species towards a common spectrum; and cohort renewal covaries with leaf lifespan, structure and physiology [Reich et al., 1991, Reich, 1995], so regularity may mark a spectrally distinctive syndrome absent from the twelve traits available to us. The crown-level analysis points to a persistent property rather than a transient state: a crown’s own regularity predicted correct classification more strongly than crown size, liana infestation or field integrity score, even though the UAV series began four years after the hyperspectral acquisition, so phase alignment in 2020–2024 need not have held in 2016.

The alternative is that a crown which does not follow its species’ cycle is simply a poorer crown. The evidence is against this as the main explanation: the field integrity score and liana presence were only modestly related to crown regularity and neither predicted classification once regularity was included, and mislabelled or contaminated crowns should inflate within-species spectral variability, which was unrelated to regularity. Field misidentification leaves no trace in either indicator, however, so a contribution from crown quality cannot be excluded; nor can a shared dependence on crown visibility, since regularity was higher in species with larger crowns (*ρ* = 0.35), although the association with F1 survived adjustment for crown area and height.

In practice, species with weak or irregular leaf cycles, including those that exchange leaves continuously, will remain harder to map from single-date imagery, and within a species the crowns that best express its leaf cycle are the ones most likely to be mapped correctly. Multi-temporal acquisitions can improve species classification [Hesketh and Sánchez-Azofeifa, 2012, Takahashi Miyoshi et al., 2020, Modzelewska et al., 2021]; our results suggest the gain would come from contrasts among species in how reflectance changes through time, rather than from capturing conspecifics in phase.

### 4.3 Sampling and signal quality: crowns, dispersion and lianas

**Number of crowns:** Within this site, approximately 17 labelled crowns per species (95% CI 14–21) were needed for an expected F1 of 0.7, and 80% of the 169 classified species had fewer (Fig. 2): label scarcity, not spectral similarity, is the first limit on species mapping in hyperdiverse forest [Baldeck and Asner, 2014, Graves et al., 2023, Mouret et al., 2024]. Improving predictions for such taxa will require larger labelled crown datasets, targeted field sampling, and emerging few-shot learning [Gevaert et al., 2025, Mu et al., 2025] and foundation-model [Huo et al., 2025] approaches.

**Spatial dispersion:** With sample size equalised, spatial dispersion of individuals across varied environmental conditions had the largest and most stable effect in the confirmatory models. Spatially clustered samples usually inflate apparent accuracy, because training and validation crowns share local conditions [Kattenborn et al., 2022]; we found the opposite, indicating that the classifier benefited from crowns sampled under diverse illumination, view geometry and site conditions, learning the species rather than the place [cf. Carlson et al., 2007, Marconi et al., 2022]. Dispersion was unrelated to any spectral metric, consistent with an effect on how representative the training sample is rather than on where the species sits in spectral space. The same reasoning bears on habitat specialists, which are the rule rather than the exception here: nearly three-quarters of the common taxa at Paracou are associated with a topographic or soil habitat [Allié et al., 2015]. A species confined to one topographic or edaphic position is sampled, unavoidably, from a narrow range of conditions, so its classifier is more likely to learn the place than the species; generalists should both classify better within a site and transfer better beyond it. *Eperua falcata*, whose trees are associated with seasonally flooded forest [Baraloto et al., 2007], was among the three least dispersed species here and among the least separable (F1 = 0.65). Dispersion reflects habitat breadth only in part, since it also depends on where the plots fall, so we offer this as a hypothesis rather than a result.

**Lianas:** Species whose crowns were more frequently infested by lianas showed reduced classification accuracy, confirmed at both the species level and the crown level. This likely reflects spectral mixing: liana leaves differ spectrally from those of their hosts [Castro-Esau et al., 2004], and infestation alters crown reflectance [van der Heijden et al., 2022], so an infested crown is a mixture, and species prone to infestation tended to be less spectrally isolated (*ρ* = −0.25, *p* = 0.09). Because liana presence was recorded opportunistically, undetected infestations may make the true effect stronger than the odds ratio of 0.44 suggests.

**Scale-dependent effects of crown size:** Crown area strongly affected whether an individual crown was classified correctly, yet mean crown area did not predict species-level F1. This discrepancy arises because crown area varies substantially within species and its effect operates at the individual level, through the number of pixels and the share of edge pixels mixed with neighbours, understorey or shadow. When mapping, classification confidence should account for crown size and visible liana contamination.

### 4.4 Traits and phylogeny: weak guides to spectral distinctiveness

Leaf chemistry and structure determine leaf reflectance, and species differ in both [Asner and Martin, 2009, Ustin et al., 2009, Asner, 2014], so it is notable that twelve leaf and bark traits told us nothing about which species were separable: this held for trait values, for trait distinctiveness, and for pairs of species, whose distance in trait space predicted neither their spectral distance nor their confusion. We see three reasons. Crown-scale reflectance depends on canopy structure, leaf angle and within-crown shading as well as on leaf optical properties, and these can dominate the signal [Ollinger, 2011, Knyazikhin et al., 2013, Townsend et al., 2013]. The traits were species means from a published dataset [Vleminckx et al., 2021], not measurements on the classified crowns, so variation among individuals and with leaf age [Chavana-Bryant et al., 2019, Bohlman, 2019] is not represented. And with 45 species we can exclude only strong trait effects. Traits surely leave their mark on spectra, but a species’ position along the measured trait axes does not predict whether it can be mapped, so accuracy is not tied to any particular functional strategy.

Phylogeny was informative only among close relatives. Reflectance showed weak phylogenetic signal across most of the spectrum (Pagel’s *λ ≈* 0.4; Fig. S21, Methods S5), in line with the phylogenetic structure reported for leaf spectra [Cavender-Bares et al., 2016, McManus et al., 2016, Meireles et al., 2020]. At the pairwise scale the structure was taxonomic rather than continuous: congeners, and to a lesser degree members of the same family, were spectrally closer and more often confused than unrelated species, whereas among species from different families relatedness made no difference. Correspondingly, species with congeners in the pool had lower F1, how close the nearest congener was did not matter, and errors in the full classification concentrated within genera. This likely reflects both spectral confusion among close relatives that share conserved leaf and crown properties and a higher risk of misidentification among congeners in the field. Accuracy will therefore depend on the composition of the species pool: forests where target species have close relatives will be harder to map, whatever their overall phylogenetic diversity.

This pattern suggested that a classifier which uses taxonomic structure might do better, as in the stepwise approach of Sapes et al. [2022]. A two-stage version suited to a multi-family community (family, then species within family; Methods S4) was less accurate than a single flat classifier (crown accuracy 0.67 vs. 0.73), because 29% of pixels were assigned to the wrong family at the first stage. Confamilial species are spectrally closer on average, but families overlap too much in spectral space for a first-stage assignment to be reliable; taxonomic information may be better used at the level of the genus, or as a penalty on taxonomically distant errors, than as a hard hierarchy.

### 4.5 Limitations and future directions

**Observational design and unmeasured confounds:** Our analysis is observational and correlational: we cannot establish that any single factor causes differences in classification accuracy. Many candidate predictors covary, and the framework does not yield causal estimates. Experimental manipulation (e.g. classifying the same species from acquisitions at different phenological stages) would be needed to isolate individual effects.

**Sample size and multiplicity:** The standardised analysis covers 45 species with complete predictor data, out of 169 classified and >800 at the site; it is biased towards common canopy species, and the results may not generalise to rare or emergent taxa, nor to sub-canopy species beyond the reach of airborne sensors. It screens 29 predictors and reports many tests without correction for multiplicity, so we rely on results that recur across methods, and note that some key coefficients, including that of regularity, are close to conventional significance thresholds.

**Unexplained variance:** About two-thirds of the cross-validated variation in species-level F1 remains unexplained. Part of this gap is noise: species-level F1 was estimated from resampled classifications of 10 crowns per species, and the uncertainty in these estimates imposes a ceiling on achievable *R*^2^. Part reflects crown-level factors that plausibly affect classification but were not measured: crown architecture and leaf angle distribution [Adams et al., 2018, Ollinger, 2011], epiphyte loading, and illumination geometry, which varies across flight lines [Arroyo-Mora et al., 2021].

**Phenology–acquisition timing mismatch:** The phenological metrics come from a 3.5-year UAV greenness series that began four years after the single hyperspectral acquisition, so the role of phenology is inferred rather than measured at the time of imaging. Greenness is an indirect proxy for leaf developmental state, and flush and shed dates derived from it are uncertain for species with complex leaf-exchange patterns. Liana presence was not surveyed systematically, and trait data were species means.

**Broader applicability:** Our findings come from a single, well-instrumented site in lowland moist forest of the Guiana Shield; the relative importance of the determinants will likely differ in forests with contrasting phylogenetic composition, phenological regimes or sensor configurations. A second hyperspectral acquisition at Paracou in 2023, contemporaneous with the phenology series, and acquisitions at two further sites in French Guiana provide the opportunity to test both the mechanisms proposed here and their generality.

### 4.6 Conclusions

This study provides the first comprehensive, multi-scale assessment of the ecological and evolutionary determinants of species-level spectral separability in a hyperdiverse tropical forest. For mapping, these results indicate that training data should favour many large, liana-free crowns sampled widely across the landscape, that classification difficulty can be anticipated from the congeneric composition of the species pool and the phenological behaviour of its species, and that single-date imaging spectroscopy will systematically favour species with regular leaf cycles.

## Acknowledgements

JGCB was supported by the NERC C-CLEAR doctoral training programme (PDAG/501), the Franklinia Foundation (through a grant awarded to DAC) and the EPIC project (Medical Research Council, MR/Z505456/1; co-PI DAC). SJ was funded by a charitable donation from John Bernstein. The work (including access to high-performance computing clusters) was supported by the Cambridge Centre for Earth Observation, directed by DAC. We thank Forestmap.ai Research for providing computational resources used in this study. Data collection in French Guiana was supported by CNES, which funded the 2016 hyperspectral data over Paracou and Labex CEBA (ANR-10-LABX-25), which funded the UAV RGB collections and the field validation of manual crown segmentations as part of the Phenobs project. Thanks to Jean-Louis Smock (IRD), Ilona Clocher (CNRS), Isabelle Maréchaux (INRAE), Chantal Geniez (IRD), Julien Engel (IRD), Tom Hattermann (CNRS), Géraldine Derroire (CIRAD), Patrick Heuret (INRAE), and all staff at Paracou Research Station for help in the field. Thanks to Philippe Verley (IRD) for support in data processing.

## Competing interests

The authors declare that they have no conflict of interest.

## Author contributions

JGCB wrote the manuscript and all authors contributed to the final version. JGCB, SJ, GV and DAC conceived the study design. JGCB and SJ ran the model training and supporting experiments. AL and CP developed the hyperspectral data from raw to analysable states. GV supervised the fieldwork and hyperspectral data collection. NB supervised the UAV data collection. AM supervised the computational infrastructure. DAC supervised the PhD of JGCB, which included contributing to the drafting of the manuscript and supervision of the analyses in Cambridge.

## Use of AI tools

AI-based tools (Claude, Anthropic) were used to assist with analysis code, to run and check supplementary analyses during revision, and to edit the text drafted by the authors. All analyses, results and interpretations were reviewed and verified by the authors, who take full responsibility for the content.

## Data availability

Hyperspectral, RGB and tree crown data will be made available upon acceptance for publication.

All classification and analysis code is in the https://github.com/PatBall1/crown-mapping repository, which will be made public upon acceptance for publication.

## Supporting Information

Methods S1–S5, Tables S1–S4, Figs S1–S21.

## Notes

https://github.com/sadiqj/hyper-analysis

### Competing Interest Statement

The authors have declared no competing interest.

### Summary of Updates

Updated analyses based on reviews. Text rewritten.

