## Supplementary Materials for "Consistent crown phenology, not functional traits, predicts whether tropical trees can be mapped from imaging spectroscopy"

### Supporting Information

#### Supplementary methods

##### Methods S1 Spectral distinctiveness metrics

**Preprocessing.** Per-pixel illumination was estimated [Schläpfer et al., 2018], and spectra were extracted from overlapping flight lines (rather than a mosaic) to exploit multiple views of each crown [Laybros et al., 2019]. Reflectance normalisation [Dalponte et al., 2014] and standard scaling were applied to all pixels. Co-registration used the LiDAR-derived canopy height model (CHM) as the baseline, with affine transformations based on eight manually assigned control points. Full processing details are in Ball et al. [2026]. The full 169-species classification used a 20-times-repeated stratified-group five-fold cross-validation over the 2,762 filtered crowns, with pixels grouped by crown so that no crown was split across folds and each crown labelled by majority vote of its pixels.

**Estimator.** Spectral metrics were computed from crown-level spectra (the per-band median of the standard-scaled pixel spectra of each crown) for the 1,969 crowns of the 45 species in the standardised dataset (10–218 crowns per species). We used crown rather than pixel spectra because the unit of classification, of the field label and of every ecological predictor is the crown; pixel-level distances would be dominated by within-crown variation (illumination, shadow, edge mixing) that the classifier averages away when pixel predictions are aggregated to a crown label. The crown spectra were projected onto their first five principal components (96.9% of variance). For each of 200 random draws of ten crowns per species, we computed the mean vector $\boldsymbol{\mu}_i$ and covariance matrix $\boldsymbol{\Sigma}_i$ of each species in this space and the Bhattacharyya distance between every pair of species,

$$D_B(i,j) = \frac{1}{8}(\boldsymbol{\mu}_i - \boldsymbol{\mu}_j)^T \boldsymbol{\Sigma}^{-1}(\boldsymbol{\mu}_i - \boldsymbol{\mu}_j) + \frac{1}{2}\ln\frac{|\boldsymbol{\Sigma}|}{|\boldsymbol{\Sigma}_i|^{1/2}|\boldsymbol{\Sigma}_j|^{1/2}}, \qquad \boldsymbol{\Sigma} = (\boldsymbol{\Sigma}_i + \boldsymbol{\Sigma}_j)/2,$$

and averaged the distances over draws. Spectral isolation of a species is its mean distance to the other 44 species, and nearest-neighbour spectral distance is its smallest distance to any other species. Within-species spectral variability is the standard deviation among the ten drawn crowns, averaged over the 378 bands and over draws.

**Why a reduced space and equal sample sizes.** With 378 bands and 10–218 crowns per species, sample covariance matrices are singular, and a Bhattacharyya distance computed from them with a small ridge added to the diagonal is governed by the number of crowns rather than by the spectra. We verified this for a full-band estimate with a ridge of $10^{-6}$: the resulting spectral isolation had a Spearman correlation of $-0.96$ with the number of crowns per species, and pairwise distance a correlation of $-0.95$ with the summed number of crowns of the two species. With five components and ten crowns per species every covariance matrix is of full rank and no ridge is needed, and equal sample sizes remove any dependence of the estimate on sampling effort: the correlation of the metrics used here with the number of crowns per species was $-0.04$ for spectral isolation, $0.05$ for nearest-neighbour distance and $0.13$ for within-species variability. For the same reason we measure within-species variability as a standard deviation rather than a coefficient of variation, which is undefined for standardised bands whose means are close to zero.

**Sensitivity analysis.** As an alternative we computed, for the same draws, the Mahalanobis distance between species means under the pooled within-species covariance matrix in the space of the first 20 principal components (99.6% of variance), the geometry in which LDA separates classes, and within-species dispersion in the same whitened space. Mahalanobis distance predicted pairwise confusion (Mantel $r = -0.34$, $p < 0.001$; $r = -0.34$ after controlling for patristic distance) about as well as Bhattacharyya distance ($r = -0.32$). At species level, Mahalanobis isolation ($\rho = 0.02$, $p = 0.90$) and nearest-neighbour distance ($\rho = 0.13$, $p = 0.38$) did not predict F1, but within-species dispersion in the whitened space did ($\rho = -0.43$, $p = 0.003$). In the Mantel tests of Table 3, the only result that differed between the two distances was that trait distance predicted Mahalanobis distance weakly ($r = 0.19$, $p = 0.03$) but not Bhattacharyya distance ($r = 0.11$, $p = 0.13$); trait distance did not predict confusion under either.

**Band-restricted metrics.** Bands were ranked by the two importance measures of Ball et al. [2026]: variance-weighted squared LDA scalings and squared structure coefficients, each averaged over 100 cross-validation folds. For the top 25% (94 bands) and top 10% (38 bands) under each ranking, the crown spectra were restricted to those bands and the whole procedure above, including the principal component projection, was repeated (Table S2). The two rankings select largely different bands: the top 10% share 5 of 38 bands.

##### Methods S2 Phylogenetic metrics

For each focal species $s$ in the species pool $S$, with $d(s,j)$ the patristic (branch-length) distance between species on the phylogeny, nearest-neighbour phylogenetic distance and mean phylogenetic distance are

$$\mathrm{NNPD}_s = \min_{j \in S \setminus \{s\}} d(s,j), \qquad \mathrm{MPD}_s = \frac{1}{|S|-1} \sum_{j \in S \setminus \{s\}} d(s,j),$$

and evolutionary distinctiveness (fair proportion) partitions the length $L_b$ of each branch $b$ on the path from $s$ to the root equally among the $N_b$ tips descended from it, $\mathrm{ED}_s = \sum_b L_b/N_b$. Nearest-congener distance is the patristic distance to the closest species of the same genus in the pool and is set to zero for species with no congener, which are also identified by a binary indicator. All metrics were computed using `ape` [Paradis and Schliep, 2019] and `picante` [Kembel et al., 2010] on a pruned `V.PhyloMaker` phylogeny [Jin and Qian, 2019]. Of the 45 species analysed, 25 had a congener in the crown dataset, and 16 had a congener among the 49 species classified together in the standardised dataset.

**Taxonomic structure of classification errors.** For a misclassified crown of species $s$ in a pool of $S$ species with $g$ congeners, the expected probability of a congeneric error under random reassignment, each other species being equally likely, is $g/(S-1)$, and likewise for confamilial errors. The ratio of observed to expected counts is reported with a 95% interval obtained as the Wilson interval on the observed proportion of misclassified crowns divided by the expected proportion.

##### Methods S3 Synchronicity, regularity and single-date homogeneity

To compare regularity and synchronicity at the level of pairs of crowns, we recomputed both for every pair of conspecific crowns in the 45 species (24,644 pairs; a random 50 crowns were used for species with more), using the daily-interpolated greenness series from October 2020 to May 2024. Species means of the recomputed metrics reproduced those used in the main analysis (Spearman $\rho = 0.98$ for regularity, 0.99 for synchronicity, 0.92 for mean absolute lag). Across all pairs, the lag at which cross-correlation peaked had a median of 53 days; it was under 30 days for 42% of pairs and over 90 days for 43%.

**Reliability.** To ask whether one metric is a less noisy species-level estimate than the other, we split the crowns of each species at random into two halves, computed both metrics within each half, and correlated the two halves across species (500 random splits). The median split-half correlation was 0.50 for regularity and 0.53 for synchronicity, corresponding to Spearman–Brown reliabilities of 0.66 and 0.69 for the full samples; regularity was the more reliable of the two in 36% of splits.

**Further phenology metrics.** Besides synchronicity, regularity and lag (main text, Methods), three descriptors of the greenness cycle entered the univariate screen and the elastic net: *amplitude*, the difference between the 99th and 1st percentiles of a crown's greenness series; *period*, the dominant cycle length from a Lomb–Scargle periodogram; and *seasonality concentration*, the Rayleigh $\rho$ of peak flush dates among conspecifics (1 when all crowns flush on the same date, 0 when flush dates are spread uniformly through the year).

**Leaf-cycle timing metrics.** From the circular mean flush and shed dates of each species we computed *leaf age at scan* (forward angular distance from mean flush date to the acquisition date, day of year 263), *leaf-age SD* (its standard deviation among conspecific crowns), and *flush* and *shed proximity* (symmetric angular distances between mean flush or shed date and the acquisition date).

**Greenness dispersion on the acquisition date.** As a direct measure of how uniform a species is on a given day, we standardised each crown's greenness series to zero mean and unit variance and took the standard deviation among conspecific crowns on 19 September of each year in the series (2021–2023), averaged over years. Using a window of $\pm 15$ days gave the same result (correlation with F1: $\rho = -0.10$, $p = 0.52$).

**Variability in the classifier's geometry.** Regularity might reduce within-species variability only in the directions that the classifier uses, leaving overall variability unchanged. We tested this with within-species dispersion measured relative to the pooled within-species covariance, the scaling that LDA applies (Methods S1). This measure predicted F1 ($\rho = -0.43$, $p = 0.003$) but was only weakly, and not significantly, related to regularity ($\rho = -0.19$, $p = 0.21$), so the data do not support this route either.

**Comparison of correlations.** The difference between the correlations of F1 with regularity and with synchronicity was tested with Williams' test for dependent correlations applied to ranks ($t = 1.16$, $p = 0.25$). Because regularity is the maximum of the cross-correlation over lags, it exceeds synchronicity even for unrelated series; this common offset does not affect which of the two predicts F1, and it does not scale with sample size (regularity versus number of crowns, $\rho = 0.13$) or differ in reliability between the metrics (above).

**Regularity and deciduousness.** To check that regularity does not simply index deciduousness, we compared the phenology metrics of the 45 species with the Paracou phenophase field record, in which trees are scored as leafy, leafless, flushing or flowering at repeated surveys (65 survey dates, October 2020 to January 2024); 31 of the 45 species have scored trees (3–45 per species, 539 in all). Regularity was unrelated to the fraction of observations in which a tree was leafless ($\rho = -0.15$, $p = 0.42$) and to the fraction of trees ever recorded leafless ($\rho = -0.22$, $p = 0.23$), whereas amplitude tracked leaf loss ($\rho = 0.50$, $p = 0.004$, with the fraction of trees ever leafless). F1 was, if anything, lower in species more often recorded leafless ($\rho = -0.38$, $p = 0.036$), and the ten species never recorded leafless had a median F1 of 0.88 against 0.82 for the rest.

**Crown-level regularity and crown quality.** The crown integrity score is a 1–4 scale of confidence that the polygon is a single, complete crown; only crowns scored 1 or 2 enter the classification, and we contrast score 1 with lower scores. Crown regularity and synchronicity were computed for the 2,294 crowns of the 45 species in the phenology series, as the mean of the pairwise metrics above over each crown's conspecifics; species means of the crown values reproduce the species-level metrics. The field database records a crown integrity score for 757 of these crowns (38% scored 2 or lower on the 1–4 scale; every crown in the classification is scored 1 or 2), and liana presence for all. A linear mixed model of crown regularity on integrity score (1 vs. lower) and liana presence with a species random intercept gave within-species effects of $-0.017$ ($t = -3.0$) for a lower integrity score and $-0.022$ ($t = -2.3$) for liana presence, against a within-species SD of crown regularity of 0.070. The crown-level GLMMs of the main text were fitted to the 535 crowns of the standardised classification for which all of crown regularity, integrity score and liana presence were available, with crown regularity and synchronicity centred within species.

#### Methods S4 Hierarchical classification test

To test whether a classifier that uses taxonomic structure improves accuracy, we compared a flat classifier with a two-stage hierarchical one on the 2,519 crowns (169 species, 41 families) of the classification training partition, using a single crown-grouped split stratified by species (1,998 training and 521 test crowns). Pixel spectra were reduced to 168 principal components (95% of variance) and classified by LDA (`MASS::lda`); crown labels were assigned by majority vote of pixels. The flat classifier predicted species directly. The hierarchical classifier predicted family first and then species with a separate LDA fitted within each of the 25 families that had more than one species, either by committing to the predicted family (hard cascade) or by weighting the within-family species posteriors by the family posterior (soft cascade). This preprocessing differs from the classifier of Ball et al. [2026], so the comparison of interest is between the flat and hierarchical variants, not with the accuracies reported there.

The flat classifier reached a crown-level accuracy of 0.73 (pixel-frequency-weighted F1 0.74, macro F1 0.46); the hard and soft cascades reached 0.67 (weighted F1 0.68, macro F1 0.36 and 0.37). The family stage assigned 71% of test pixels to the correct family. The change in per-species F1 relative to the flat classifier did not depend on whether a species had congeners in the pool (mean change $-0.08$ with congeners and $-0.09$ without for the hard cascade; Spearman correlation with number of congeners $-0.03$, $p = 0.74$). This is an exploratory test on a single split and does not exclude other ways of using taxonomic information.

#### Methods S5 Phylogenetic signal in reflectance

We tested for phylogenetic signal in species-mean reflectance in each of the 378 bands, for the 141 species on the phylogeny with at least 100 pixels, using Pagel's $\lambda$ (likelihood-ratio test against $\lambda = 0$) and Blomberg's $K$ (1,000 randomisations) in `phytools` [Revell, 2012]. We also tested whether F1 itself showed phylogenetic signal.

At the level of individual bands, species-mean reflectance showed statistically detectable but weak phylogenetic signal (Fig. S21). Pagel's $\lambda$ differed from zero in 302 of 378 bands (80%; median $\lambda = 0.39$, interquartile range 0.31–0.45), including all eight far-red-edge bands (median 0.50), but was indistinguishable from zero across much of the near-infrared plateau (935–1,340 nm). Blomberg's $K$ was very low throughout (median 0.021, interquartile range 0.018–0.023), as expected for a $\lambda$ of this size on a phylogeny with many very short terminal branches. Related species therefore resemble one another spectrally more than random pairs do, but much less than their shared ancestry alone would predict.

#### Supplementary tables and figures

**Table S1:** The ten UAV RGB surveys used for crown delineation [Ball et al., 2026] and as the start of the phenology series, with the aircraft used on each date.

| Survey | Date | Aircraft |
|---|---|---|
| 1 | 23-Oct-2020 | DJI Phantom 4 Advanced |
| 2 | 05-Nov-2020 | DJI Phantom 4 Advanced |
| 3 | 23-Nov-2020 | DJI Phantom 4 Advanced |
| 4 | 14-Dec-2020 | DJI Phantom 4 Advanced |
| 5 | 05-Jan-2021 | DJI Phantom 4 Advanced |
| 6 | 18-Jan-2021 | DJI Phantom 4 Advanced |
| 7 | 08-Feb-2021 | DJI Phantom 4 Advanced |
| 8 | 03-Mar-2021 | DJI P4 Multispectral |
| 9 | 16-Mar-2021 | DJI P4 Multispectral |
| 10 | 06-Apr-2021 | DJI P4 Multispectral |

**Table S2:** Spectral distinctiveness metrics computed from the full spectrum and from subsets of the most important bands (Methods S1). Columns give the Spearman correlation ($p$-value) of species-level F1 with spectral isolation, nearest-neighbour spectral distance and within-species spectral variability ($n = 45$ species), and the Mantel correlation between pairwise spectral distance and pairwise confusion (990 pairs). FRE: number of the eight far-red-edge bands (748–775 nm) included.

| Band set | Bands | FRE | Isolation | Nearest neighbour | Variability | Confusion (Mantel) |
|---|---|---|---|---|---|---|
| All bands | 378 | 8 | 0.45 (0.002) | 0.38 (0.010) | −0.31 (0.038) | −0.32 (<0.001) |
| Top 25%, LDA scalings | 94 | 8 | 0.38 (0.009) | 0.34 (0.021) | −0.33 (0.027) | −0.30 (<0.001) |
| Top 10%, LDA scalings | 38 | 8 | 0.41 (0.006) | 0.38 (0.010) | −0.35 (0.020) | −0.30 (<0.001) |
| Top 25%, structure coefficients | 94 | 0 | 0.38 (0.011) | 0.30 (0.045) | −0.40 (0.007) | −0.34 (<0.001) |
| Top 10%, structure coefficients | 38 | 0 | 0.37 (0.012) | 0.30 (0.046) | −0.37 (0.012) | −0.30 (<0.001) |

**Table S3:** Elastic-net coefficients (logit F1, standardised predictors) with the 12 leaf and bark traits entered as raw values, as in the pre-specified screen, or replaced by the two trait-distinctiveness metrics ($n = 45$ species). Both fits are from the same sensitivity run, with the mixing parameter and penalty chosen by 10-fold cross-validation ($\alpha = 0.4$ and 0.3). The raw-trait fit is a re-fit of the screen in Fig. 4 with its own cross-validated penalty, so its coefficients differ slightly and it additionally retains period with a small coefficient. Predictors not listed had coefficients of zero in both fits; a dash marks a predictor not in that pool.

| Predictor | Traits as 12 values (29 predictors) | Traits as distinctiveness (19 predictors) |
|---|---|---|
| No congener in dataset | +0.17 | +0.20 |
| Spatial dispersion | +0.13 | +0.15 |
| Mean phylogenetic distance | +0.15 | +0.14 |
| Phenological regularity | +0.12 | +0.14 |
| Liana prevalence | −0.10 | −0.12 |
| Seasonality concentration | −0.04 | −0.06 |
| Period | −0.02 | −0.03 |
| Leaf carbon | +0.07 | — |
| Trunk bark thickness | +0.03 | — |
| Trait isolation | — | −0.04 |
| Nearest-neighbour trait distance | — | 0.00 |

**Table S4:** Spearman correlations between species-level F1 and the 12 leaf and bark traits (species means from Vleminckx et al. 2021) and the two trait-distinctiveness metrics ($n = 45$ species; unadjusted $p$-values). No correlation is significant, with or without correction for multiple tests.

| Trait | $\rho$ | $p$ |
|---|---|---|
| Leaf carbon | +0.24 | 0.11 |
| Trunk bark thickness | +0.19 | 0.22 |
| Leaf thickness | −0.17 | 0.27 |
| Leaf potassium | −0.16 | 0.30 |
| Leaf toughness | −0.11 | 0.46 |
| Leaf nitrogen | +0.05 | 0.77 |
| Leaf phosphorus | +0.04 | 0.78 |
| Specific leaf area | +0.04 | 0.78 |
| Chlorophyll content | +0.03 | 0.84 |
| Leaf calcium | +0.03 | 0.86 |
| Leaf $\delta^{13}$C | −0.02 | 0.90 |
| Leaf area | −0.01 | 0.93 |
| Nearest-neighbour trait distance | +0.05 | 0.73 |
| Trait isolation | −0.23 | 0.12 |

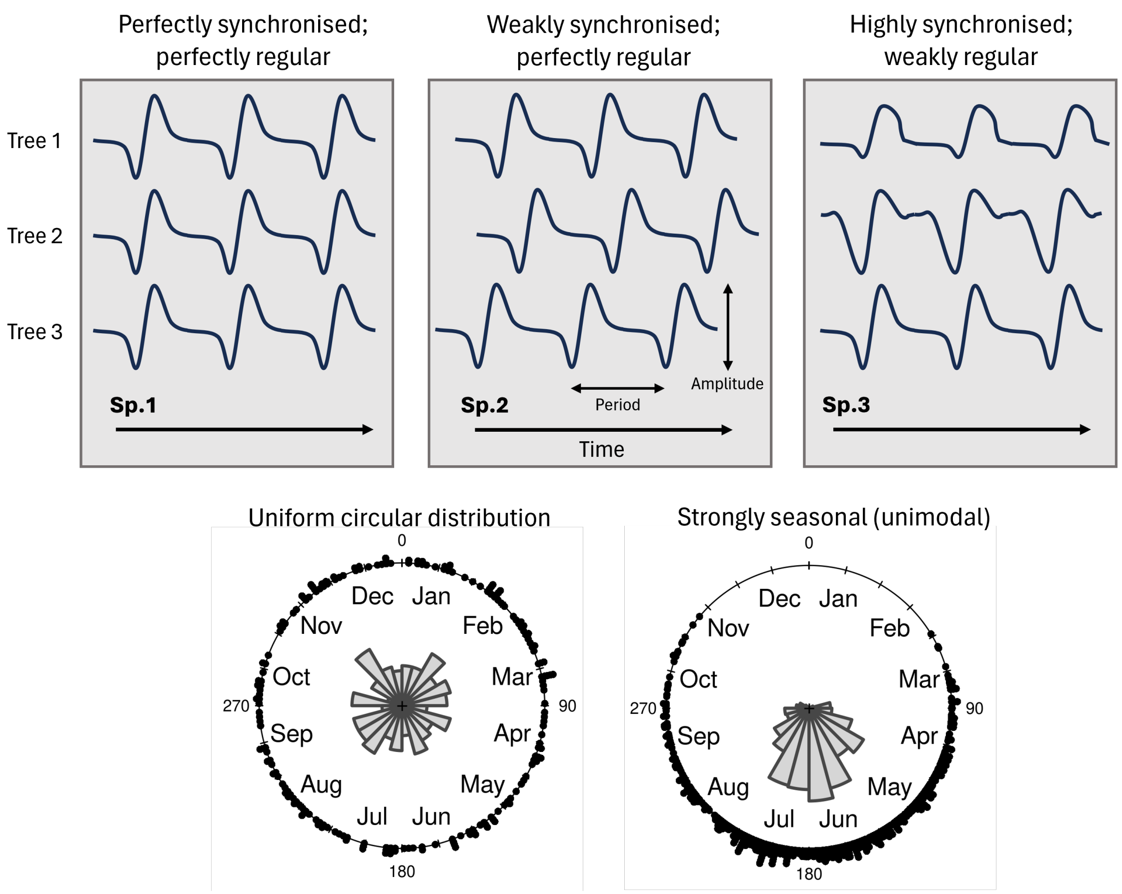


**Figure S1:** Schematic of the phenological metrics (main text, Methods). Top row: as a tree cycles through its leaves, crown greenness (GCC) oscillates with a characteristic *period* and *amplitude*. Conspecific trees may be synchronous and regular (left: waveforms aligned in time), regular but weakly synchronous (centre: the same waveform offset in time, so the cross-correlation peaks at a non-zero *lag*), or synchronous but weakly regular (right: peaks aligned but waveforms differ). *Regularity* is the maximum cross-correlation between two conspecific greenness series over lags of up to ±365 days; *synchronicity* is their correlation at lag zero; both are averaged over conspecific pairs. Bottom row: peak flush dates among conspecifics may be spread through the year (left; low *seasonality concentration*, Rayleigh $\rho$) or concentrated in one season (right; high Rayleigh $\rho$). Schematic adapted from Ball et al. (in preparation) on crown phenology at this site.

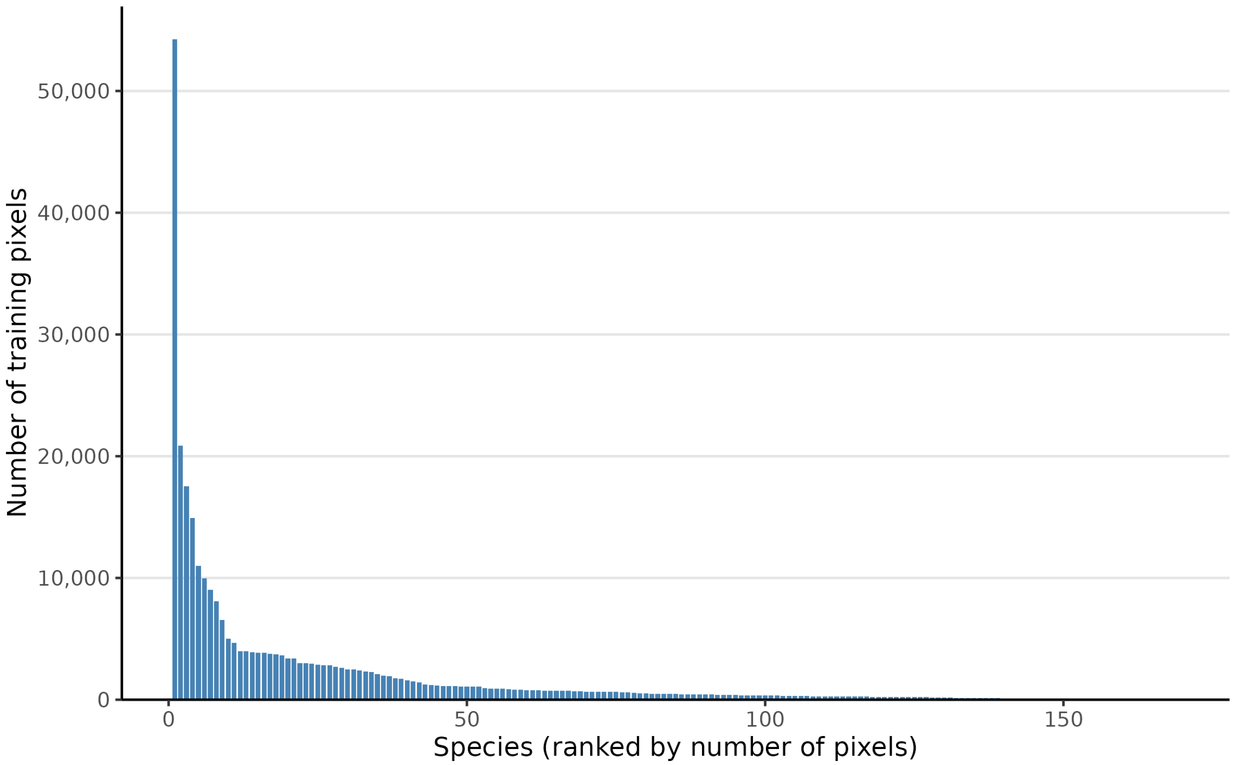


**Figure S2:** Number of training pixels per species (169 species), ordered by prevalence (1 = most prevalent). The ten most prevalent species hold 53% of the pixels.

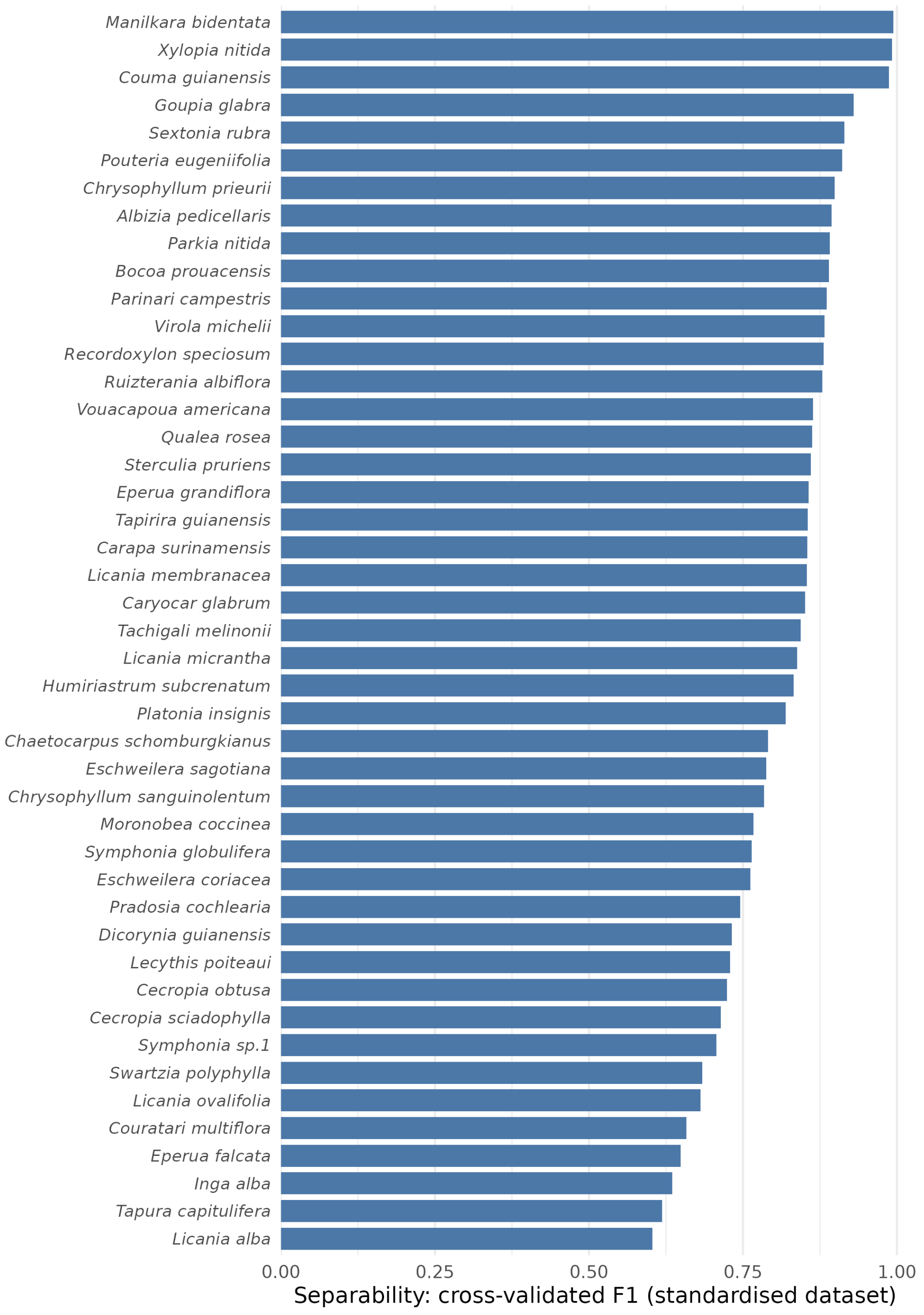


**Figure S3:** Separability (cross-validated F1) in the standardised dataset (ten crowns per species) for the 45 species analysed, ordered by F1.

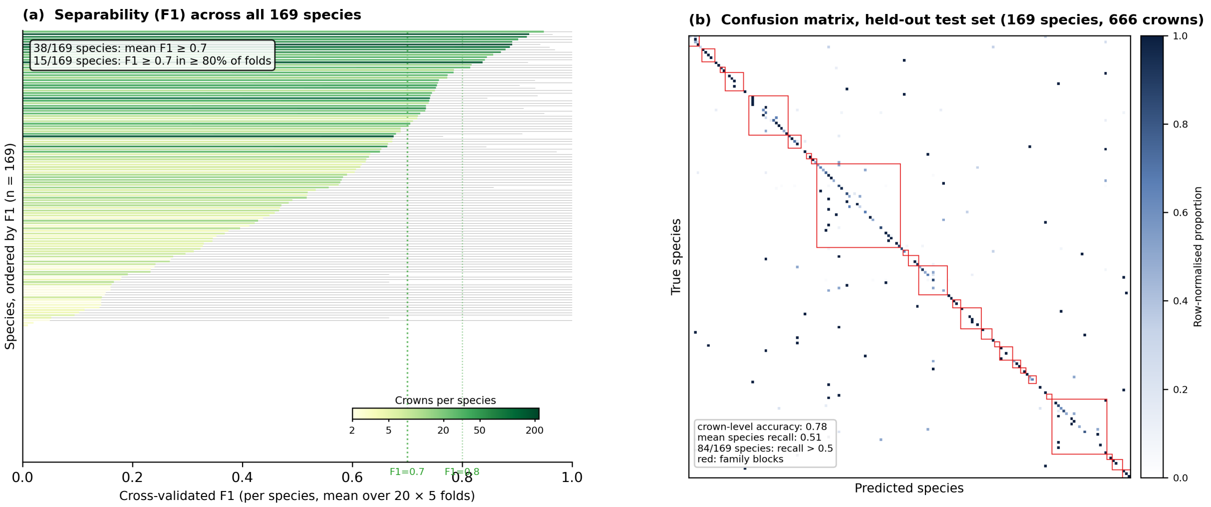


**Figure S4:** Per-species separability and the full confusion matrix. (a) Per-species mean F1 across all 169 species in the repeated 20 × 5-fold cross-validation, ordered by F1; bar colour shows the number of labelled crowns of each species on a log scale (darker green = more crowns); vertical reference lines mark F1 = 0.7 and F1 = 0.8; thin bars span the 2.5–97.5% quantiles across folds. Annotation reports how many species sit above F1 = 0.7 on average and how many do so reliably ($\geq$ 80% of folds). (b) Row-normalised confusion matrix on the held-out test set (169 species, 666 crowns; same train/test split as Ball et al. 2026). Species are sorted by family; red squares enclose multi-species family blocks; the annotation gives the crown-level accuracy and the mean per-species recall.

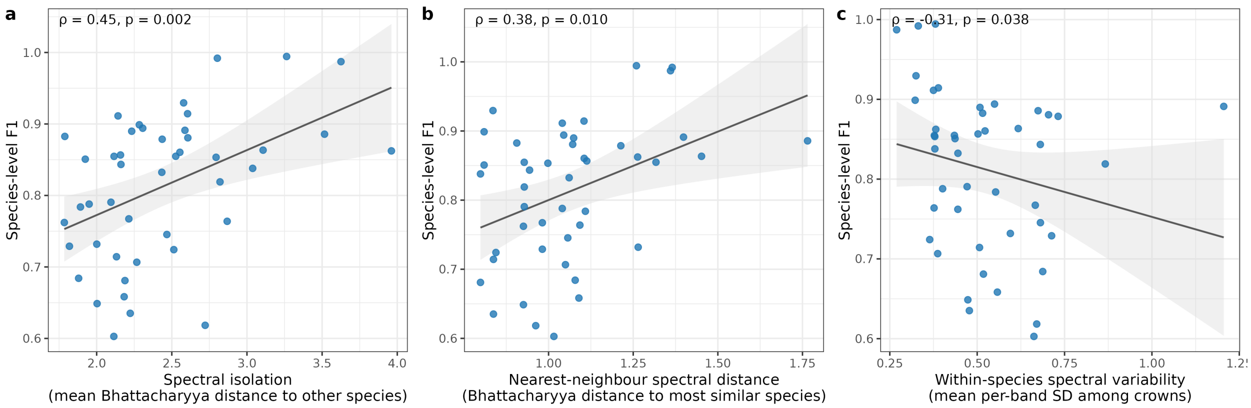


**Figure S5:** Species-level F1 versus spectral distinctiveness ($n = 45$ species; Spearman $\rho$ and $p$ on each panel; lines are linear fits with 95% confidence bands). (a) Spectral isolation, the mean Bhattacharyya distance from a species to all others. (b) Nearest-neighbour spectral distance. (c) Within-species spectral variability, the standard deviation among conspecific crowns averaged over bands. All metrics are computed from ten crowns per species (Methods S1).

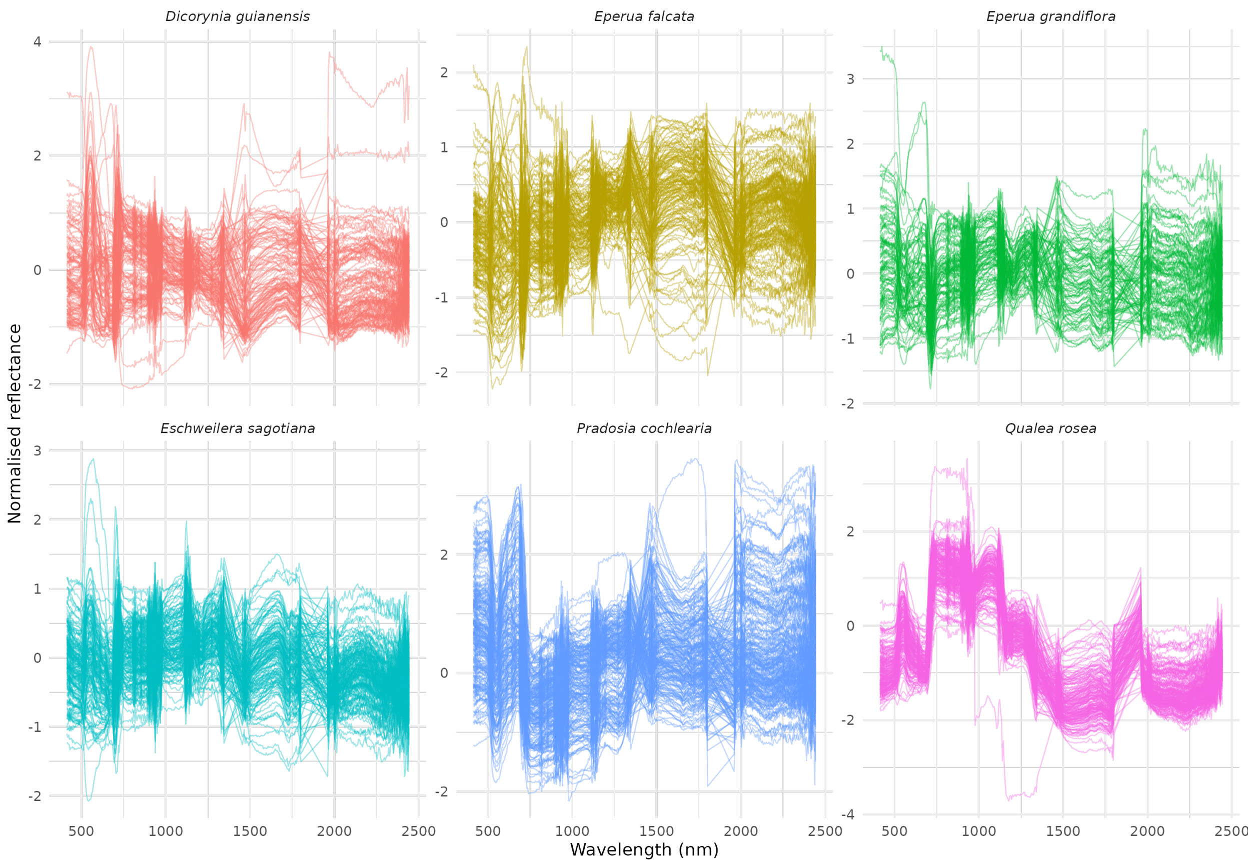


**Figure S6:** Within-species spectral variability for selected species. Each line represents the spectrum of a single crown; colour indicates species identity.

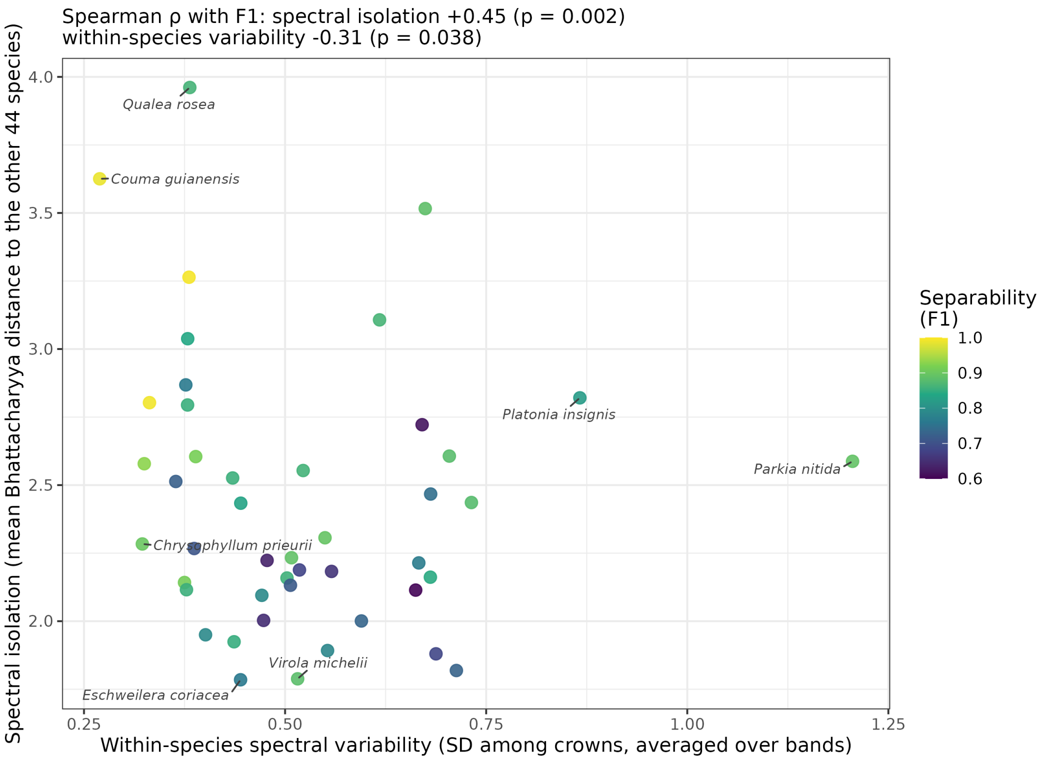


**Figure S7:** Within-species spectral variability versus spectral isolation for the 45 species of the standardised dataset (metrics as in Methods S1; each point is a species, coloured by its F1). Species that were spectrally isolated and internally consistent (upper left) tended to be more separable (Spearman $\rho$ with F1: spectral isolation $+0.45$, within-species variability $-0.31$; Fig. S5). The two metrics are nearly uncorrelated ($\rho = -0.19$), so they describe different aspects of distinctiveness. The two species at each extreme of each axis are labelled.

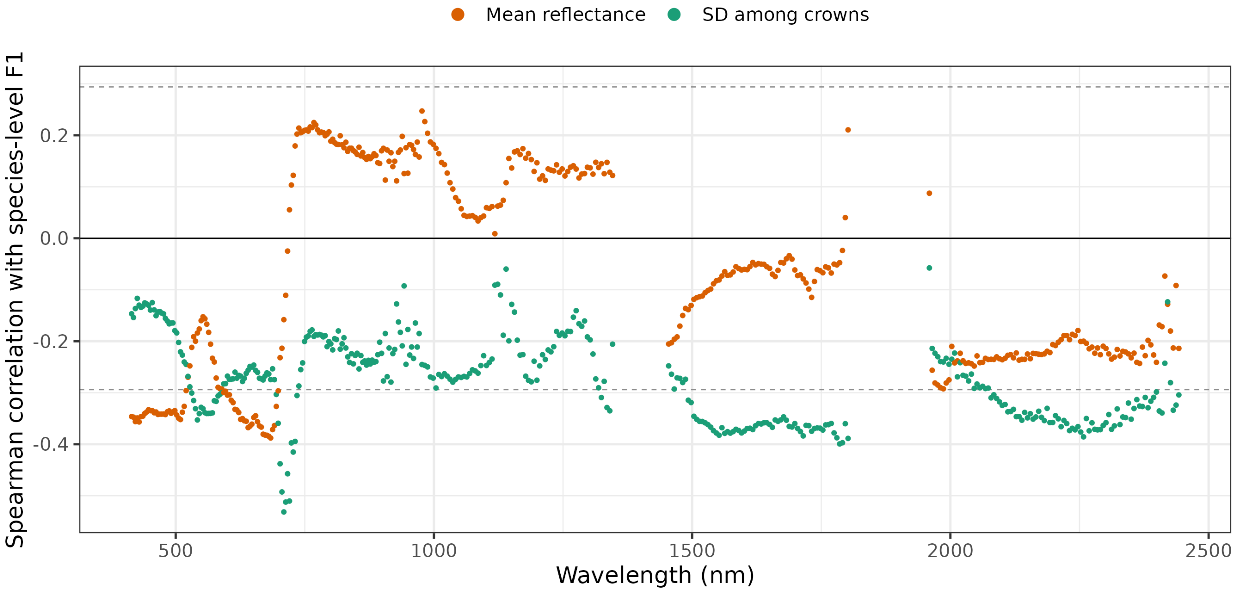


**Figure S8:** Spearman correlation between species-level F1 and, for each band, the mean reflectance of a species (orange) and the standard deviation among its crowns (green), for the 45 species of the standardised dataset. Dashed lines mark $|\rho| = 0.29$, the value at which a single correlation reaches $p = 0.05$ (uncorrected).

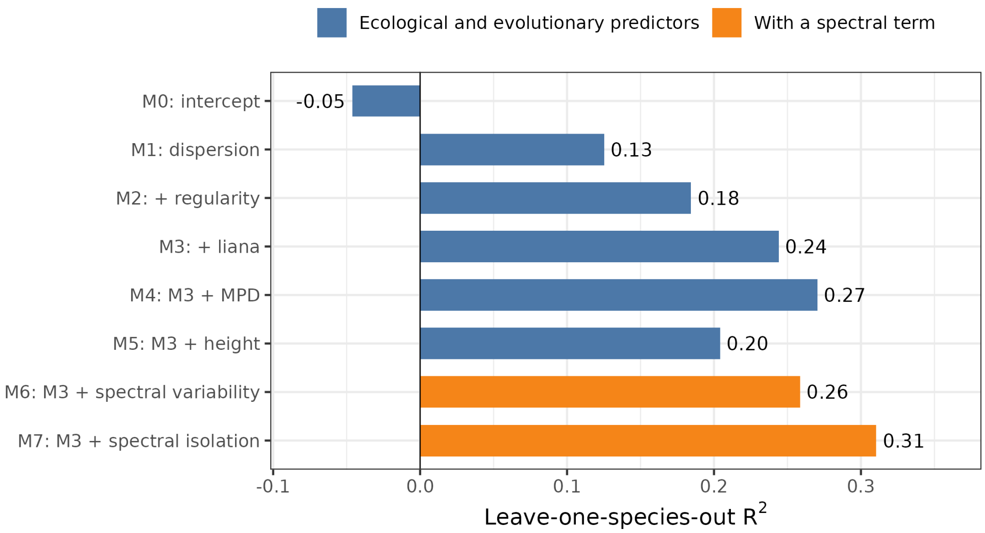


**Figure S9:** Leave-one-species-out (LOSO) cross-validated $R^2$ of the beta regression models. M0 is the intercept-only model; M1 adds spatial dispersion, M2 regularity and M3 liana prevalence; M4–M7 each add one term to M3. A model adding both spectral terms to M3 reached LOSO $R^2 = 0.33$.

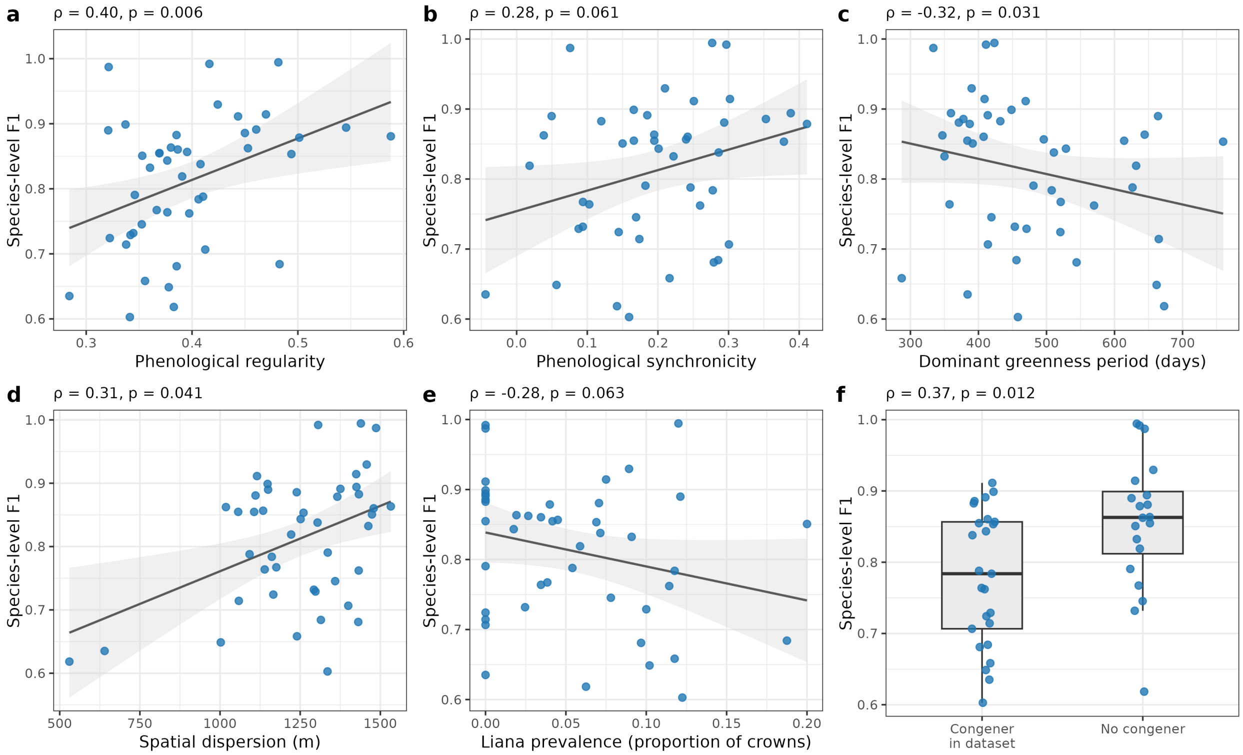


**Figure S10:** Univariate relationships between species-level F1 and six predictors ($n = 45$ species; Spearman $\rho$ and $p$ above each panel; in (a–e) lines are linear fits with 95% confidence bands). (a) Phenological regularity. (b) Phenological synchronicity. (c) Dominant period of the greenness cycle. (d) Spatial dispersion. (e) Liana prevalence. (f) Presence of a congener in the crown dataset.

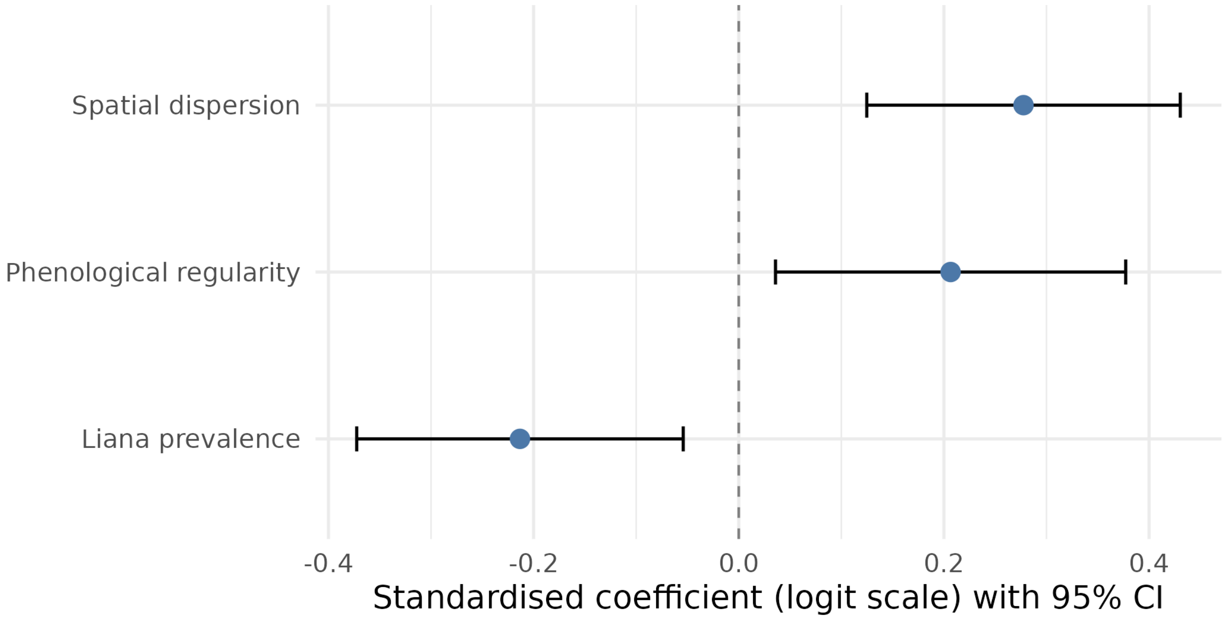


**Figure S11:** Beta regression coefficients for model M3 (spatial dispersion + phenological regularity + liana prevalence; LOSO $R^2 = 0.24$). Points show standardised coefficients on the logit scale, with 95% confidence intervals.

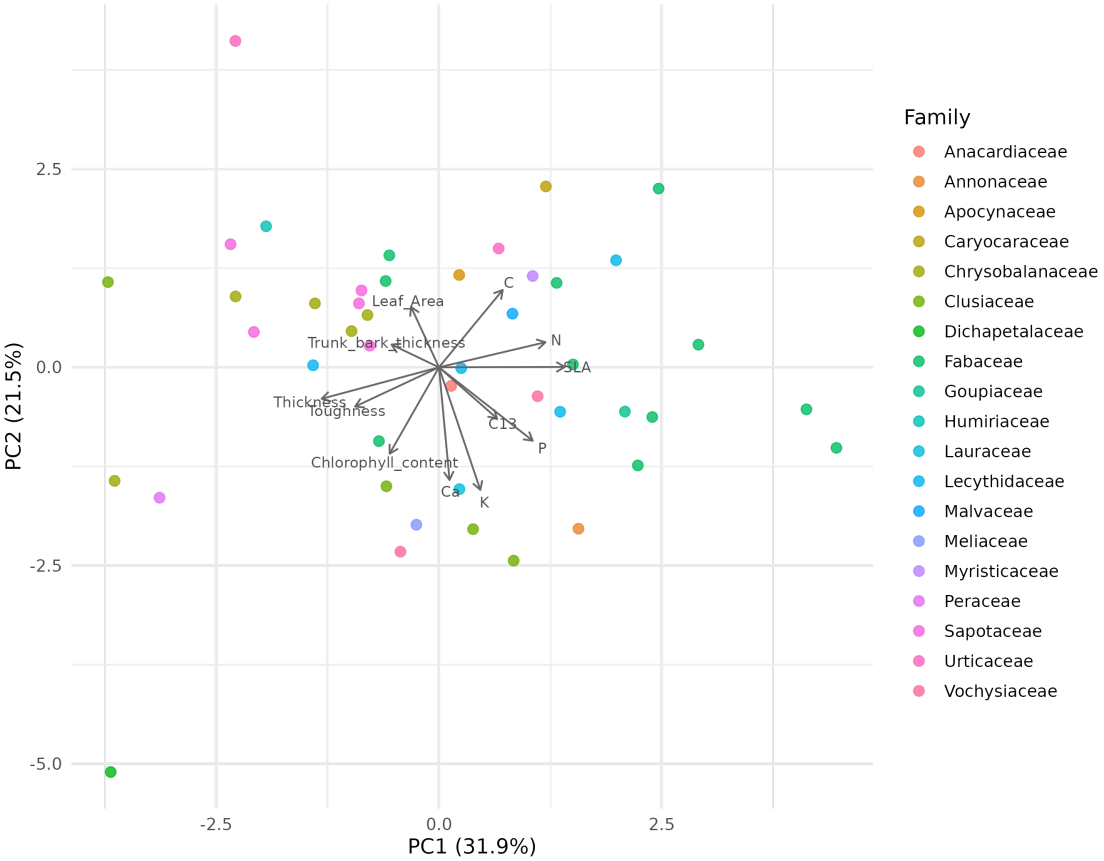


**Figure S12:** PCA of species-mean leaf and bark traits (12 traits from Vleminckx et al. 2021). Points are coloured by family.

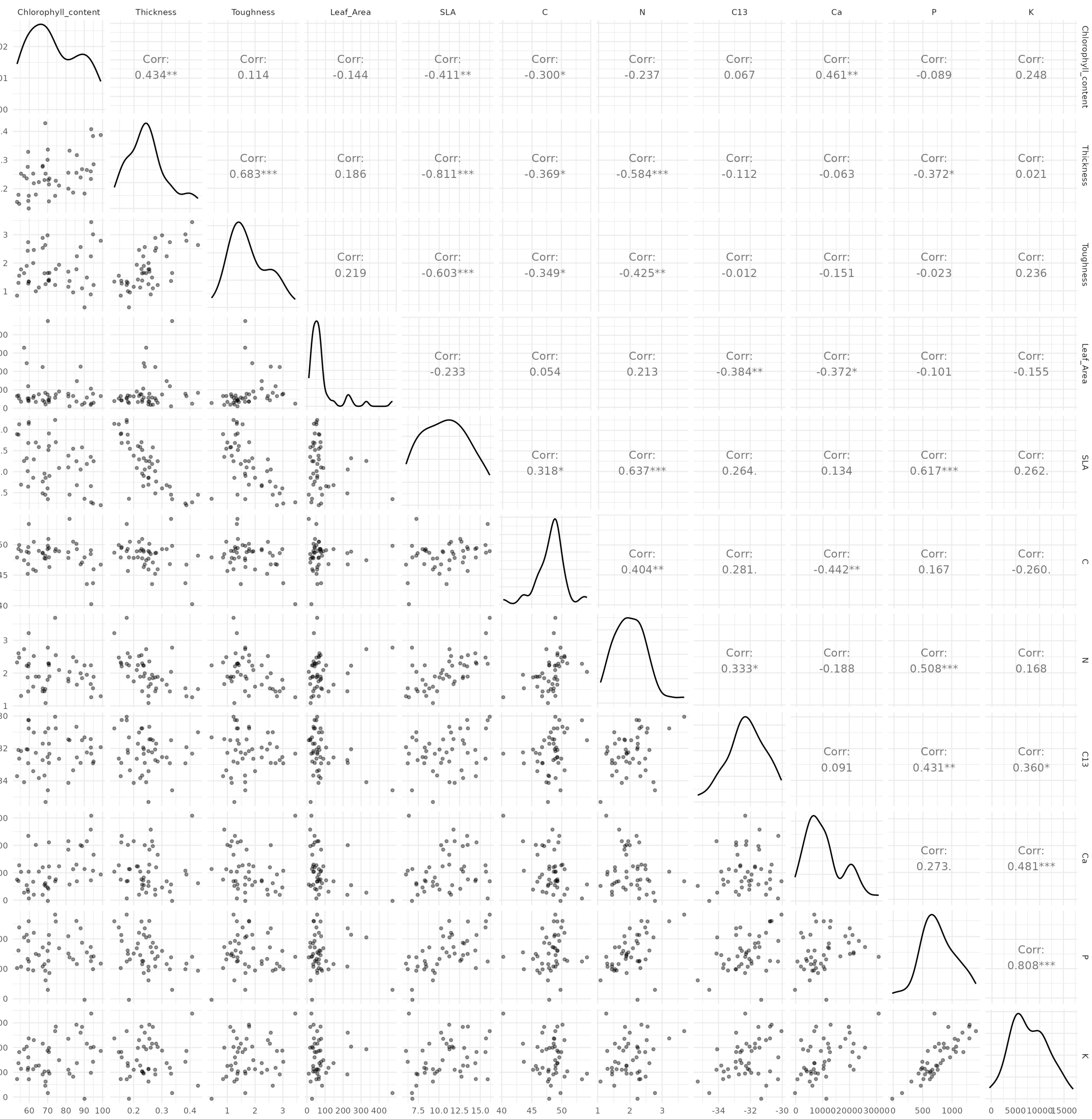


**Figure S13:** Pairwise correlations among the 11 leaf traits (trunk bark thickness is not shown; variable names as in the analysis data, e.g. C13 = $\delta^{13}$C). Upper triangle: Pearson $r$ with significance marks (*** $p < 0.001$, ** $p < 0.01$, * $p < 0.05$, . $p < 0.1$); lower triangle: scatter plots; diagonal: trait distributions.

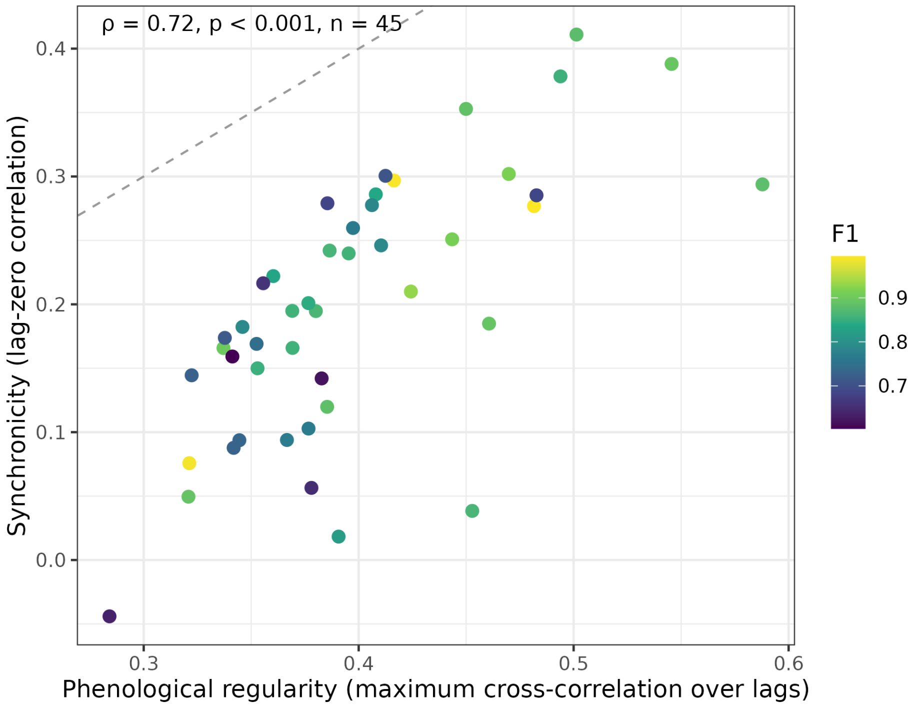


**Figure S14:** Phenological regularity versus synchronicity ($n = 45$ species; $\rho = 0.72$, $p < 0.001$). Points are coloured by species-level F1; the dashed line marks equality, which bounds synchronicity from above.

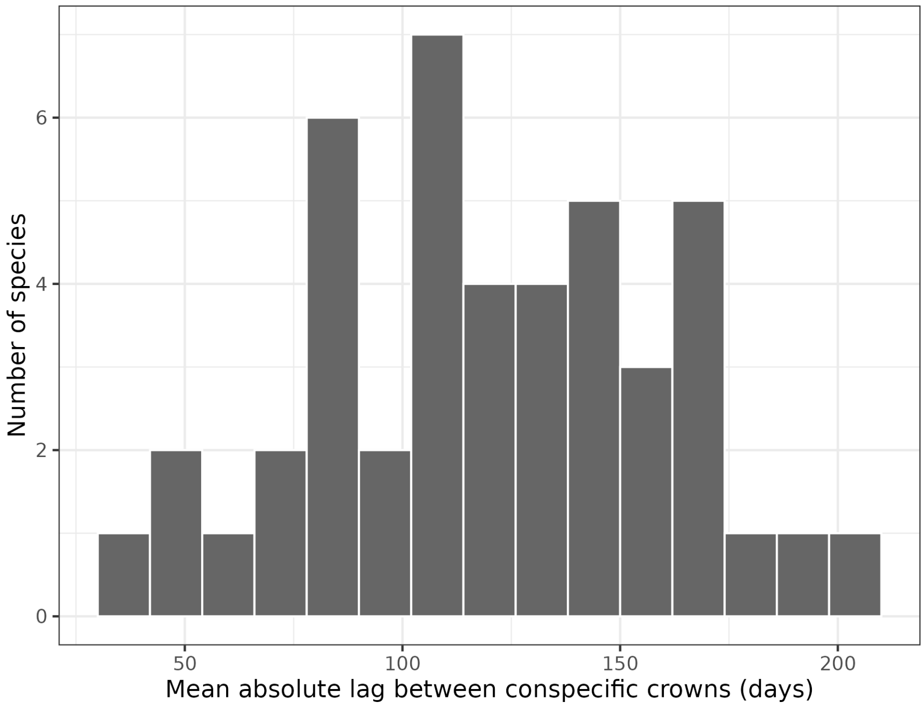


**Figure S15:** Distribution across species of the mean absolute lag (days) at which the cross-correlation between conspecific greenness series peaks ($n = 45$ species; mean 120, median 121, range 39–202 days). Lag was not associated with species-level F1 ($\rho = -0.22$, $p = 0.16$).

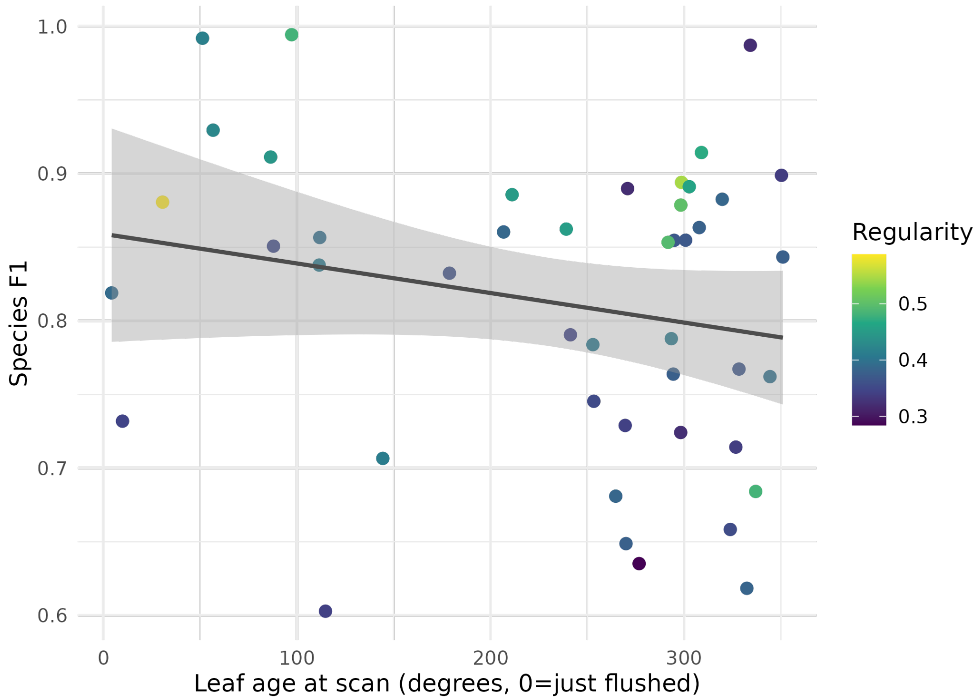


**Figure S16:** Leaf age at scan (forward angular distance from mean flush date to acquisition date, in degrees of the annual cycle) versus species-level F1 ($\rho = -0.12$, $p = 0.42$); points are coloured by phenological regularity.

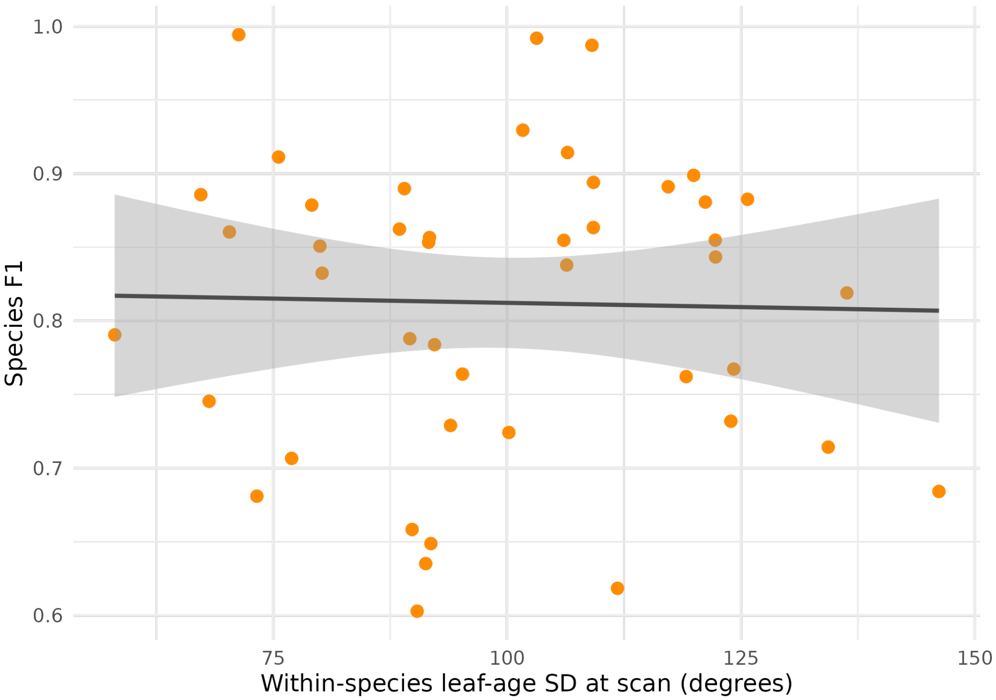


**Figure S17:** Leaf-age SD (standard deviation of leaf age at scan among conspecific crowns) versus species-level F1 ($\rho = -0.01$, $p = 0.96$).

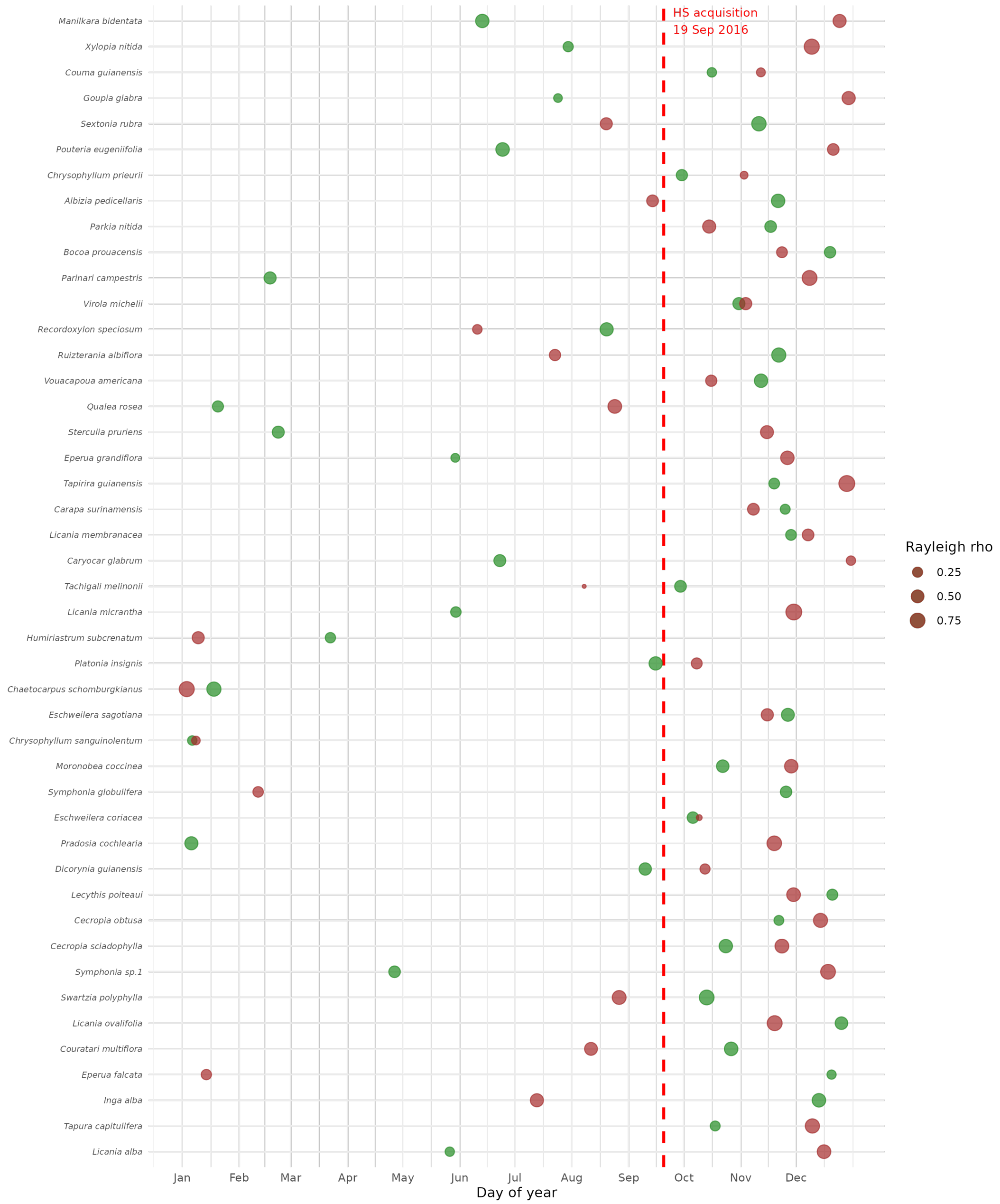


**Figure S18:** Leaf-cycle timing relative to the hyperspectral acquisition date (19 September 2016; vertical dashed line). Species are ordered by F1 (bottom = lowest). Green and red-brown points indicate mean circular flush and shed dates estimated from crown-level greenness series, respectively; point size reflects the consistency of timing among conspecifics (Rayleigh $\rho$). Flush and shed dates are turning points of the greenness series rather than observed phenophases; in species whose dominant period exceeds a year (e.g. *Platonia insignis*, 632 days) a flush and a shed date can fall close together. No timing metric predicted species separability (Figs. S16 and S17).

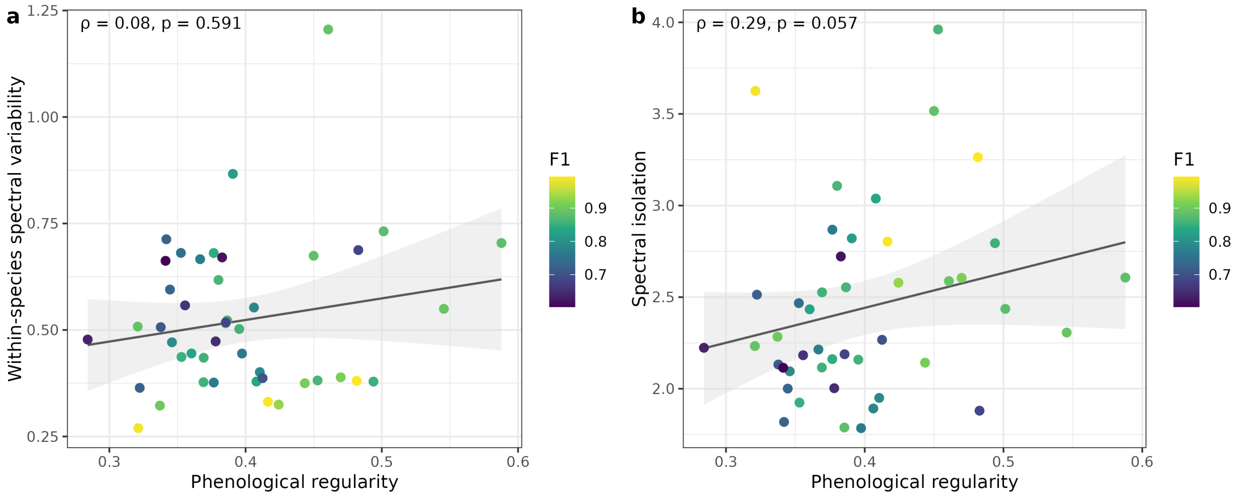


**Figure S19:** Phenological regularity and spectral distinctiveness ($n = 45$ species; points coloured by F1; Spearman $\rho$ and $p$ on each panel). (a) Within-species spectral variability is unrelated to regularity. (b) Spectral isolation tends to increase with regularity.

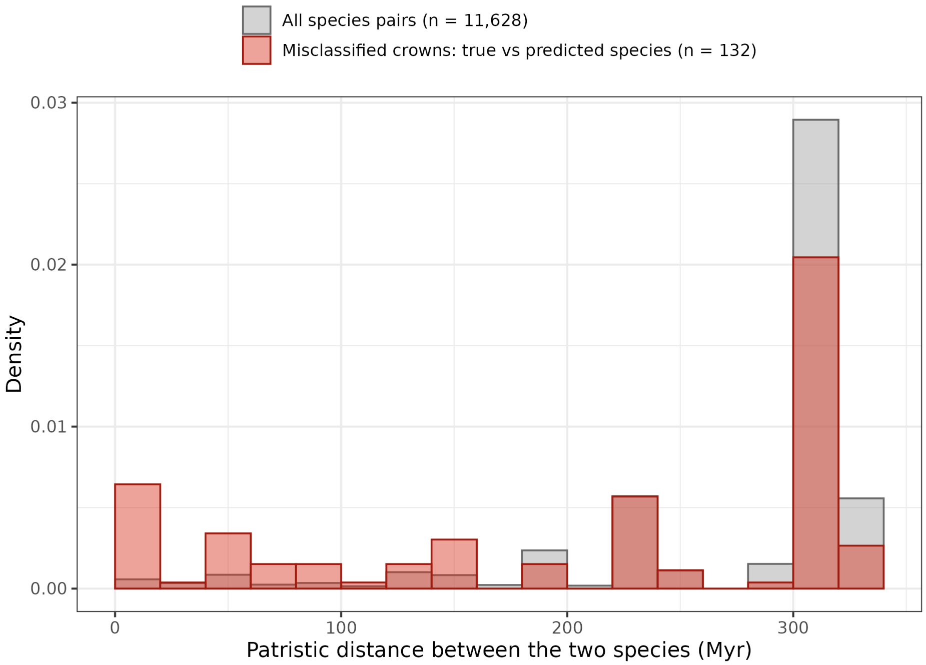


**Figure S20:** Phylogenetic distance and misclassification in the full classification. Patristic distance between the true and the predicted species for the 132 misclassified test crowns whose species are both on the phylogeny (red), compared with all 11,628 pairs among the 153 test species on the phylogeny (grey). Misclassifications fall disproportionately between close relatives: 27% lie at distances below 100 Myr, against 5% of all pairs (Wilcoxon $p < 0.001$). Across species pairs the misclassification rate declines weakly with patristic distance (Spearman $\rho = -0.05$, $p < 0.001$).

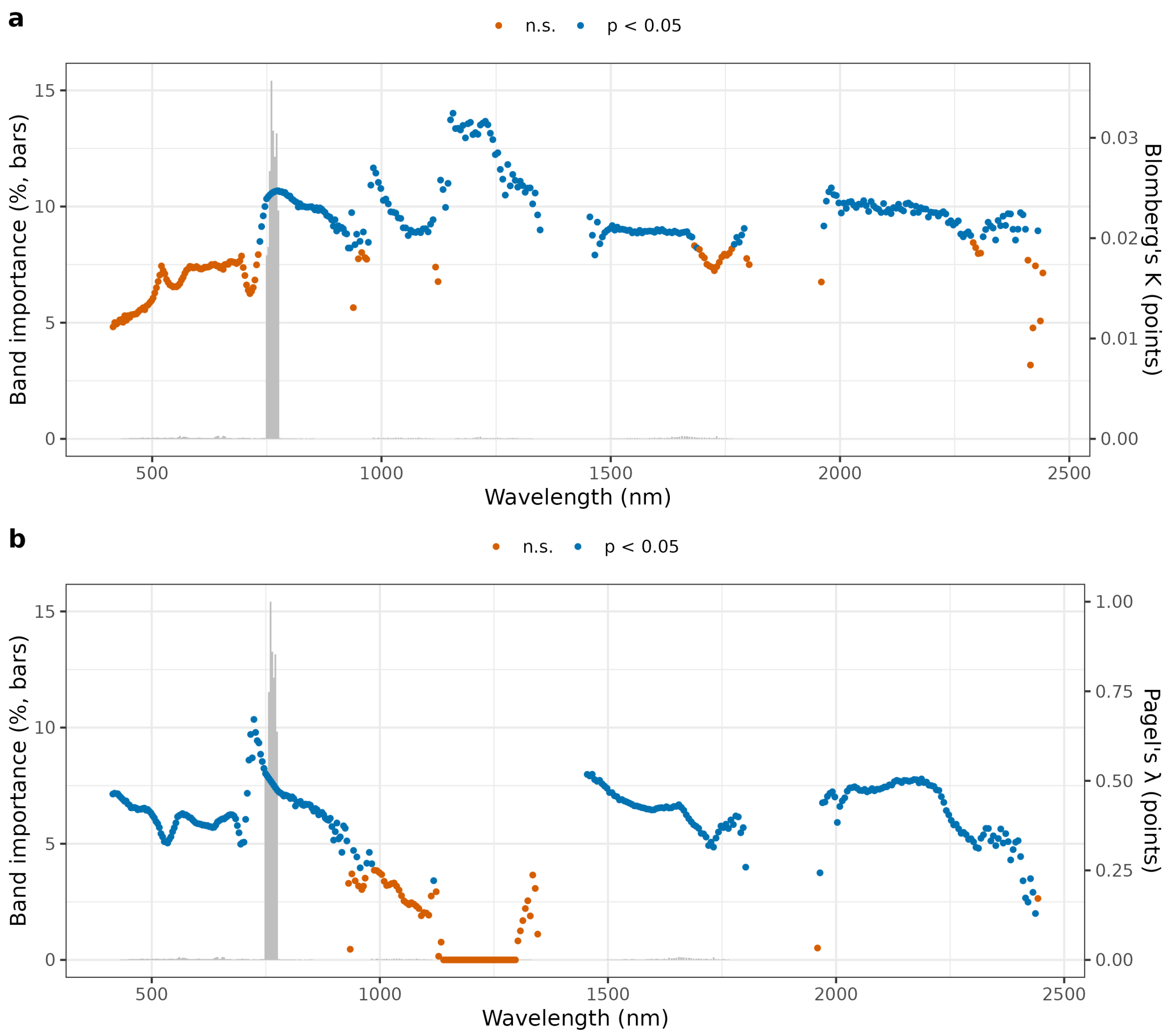


**Figure S21:** Phylogenetic signal in species-mean reflectance for each of the 378 bands (141 species). (a) Blomberg's $K$ and (b) Pagel's $\lambda$ (points, right axes), with the band importance of the classifier (variance-weighted LDA scalings, as a percentage of the total; grey bars, left axes). Blue points are significant at $p < 0.05$ (randomisation test for $K$; likelihood-ratio test against $\lambda = 0$).
